# Microenvironment optimization enables kidney organoid longevity with epithelial-endothelial joint basement membrane formation

**DOI:** 10.64898/2026.04.09.717566

**Authors:** Sophie M. Blackburn, Benjamin A. Juliar, Arjun Sen, Mary C. Regier, Benjamin S. Freedman

**Author notes:** Correspondence: Benjamin S. Freedman, Ph.D., Associate Professor, University of Washington School of Medicine, 850 Republican St, Box 358056, Seattle, WA 98109, USA. All authors agree the author list is complete and acceptable, and that the author list and order is subject to change significantly in any final, peer-reviewed publication.

## Abstract

Kidney organoids degrade in long-term culture and lack joint basement membranes between epithelial and endothelial cells characteristic of renal tissue. Here we show that these limitations can be overcome in static cultures simply by optimizing the microenvironment. Supplementing standard media with tubular-enhancing factors (TEFs) dramatically improves organoid yield and longevity, while vascular-enhancing factors (VEFs) and replating increases endothelial cell yield and invasiveness. A transcriptomic and imaging atlas demonstrates maintenance of nephron structures for six months with increased metabolism, signaling, differentiation, and aging-related pathways. In addition to adherent cultures, these media also enable organoid differentiation and vascularization in suspension cultures and hydrogels. Remarkably, addition of TEFs and VEFs to organoids in suspension induces self-assembly of joint basement membranes between endothelial cells and podocytes or tubules, a major feature of renal tissue. Microenvironment optimization thus enables longitudinal stabilization and higher-order vascularization of kidney organoids, offering a diverse resource for long-term studies and tissue engineering applications.

---

Organoids derived from human pluripotent stem cells (hPSCs) have emerged as valuable tools for disease modeling and regenerative medicine, but still face significant limitations.^1^ One major challenge is limited longevity. hPSCs typically undergo a single wave of differentiation, and the resultant organoids tend to degrade over time, undergoing epithelial dedifferentiation, stromal overgrowth, and internal necrosis.^2–5^ In a parallel system, that of intestinal organoids (enteroids), identification of a microenvironment that supports intestinal stem cells, including specialized media and three-dimensional culture, has enabled extensive propagation.^1,6^ A variation of this recipe, called tubuloid expansion media (TEM), has enabled the passaging and culture of adult kidney tubuloids for up to six months.^4,7^ In contrast to enteroids, tubuloids are not known to contain specialized adult stem cells. TEM incorporates key components such as R-Spondin 3 conditioned medium (RSPO3), epidermal growth factor (EGF), fibroblast growth factor 10 (FGF-10), and A83-01.^4,7^ Together, these factors stabilize renal tubular cells over longer periods than standard media. Recent work has used TEM to propagate tubuloids derived from hPSCs for up to three months, although whether this would stabilize organoids in the absence of passaging is not yet known.^4^ Being able to maintain organoid for longer periods would enable diverse applications, for instance studying tissue aging over time leading to chronic kidney disease, or serving as avatars for human passengers on a space flight to Mars, a journey that would require at least six months to complete.

Another area where organoid cultures need improvement is vascularization, which remains primitive even when vascular cells are present. For instance, joint basement membranes (JBMs) are highly characteristic of renal tissue *in vivo,* including both glomerular basement membranes (GBMs) and tubular basement membranes (TBMs), which perform essential physiological functions such as filtration and transport with the blood circulation.^8–10^ Kidney organoids derived from hPSCs naturally contain endothelial cells, but these fail to naturally form GBMs with neighboring podocytes or TBMs with proximal tubular cells.^11,12^ Placing organoids in flow channels or treating them with vascular endothelial growth factor (VEGF) can increase endothelial cell number and interactions with epithelial cells, but is insufficient to induce the formation of JBMs.^13–15^ Similarly, top-down bioengineering approaches can create parallel channels for seeding with endothelial and epithelial cells, but these are generally unable to bring the cells into direct contact, and the dissociation required for seeding can injure cells^16–18^.

Interestingly, implantation of organoids *in vivo* beneath the kidney capsule results in chimeric glomerulus-like structures that appear to contain GBMs.^19–21^ Various factors enabling JBM formation are therefore hypothesized to be missing in the microenvironment *in vitro,* including blood flow, cell maturation, vascular growth factors, and developmental time.

Here, to address these limitations, we investigate optimized media formulations that incorporate tubular-enhancing factors (TEFs) and vascular-enhancing factors (VEFs) in kidney organoids. Surprisingly, our studies reveal that supplementing standard media with TEFs and VEFs can have a dramatic impact on organoids in static cultures, including prolonging their survival for up to six months, and enabling the self-organization of both GBMs and TBMs.

## RESULTS

### TEFs improve kidney organoid differentiation and survival

We tested the effect of supplementing our standard organoid growth medium (RB) with tubuloid expansion medium (TEM) in a 1:5 ratio, a formulation we called OTEM (organoid tubular enhancing medium, **Supplementary Table 1**).^4,7,22^ OTEM was substituted for RB on day 11 after hPSC seeding, a time point at which renal vesicle-like structures have formed (**Figure 1A**). In both OTEM and RB, these structures proceeded to differentiate into tubular organoids along a similar time course (**Figure 1B**).

**Figure 1.**
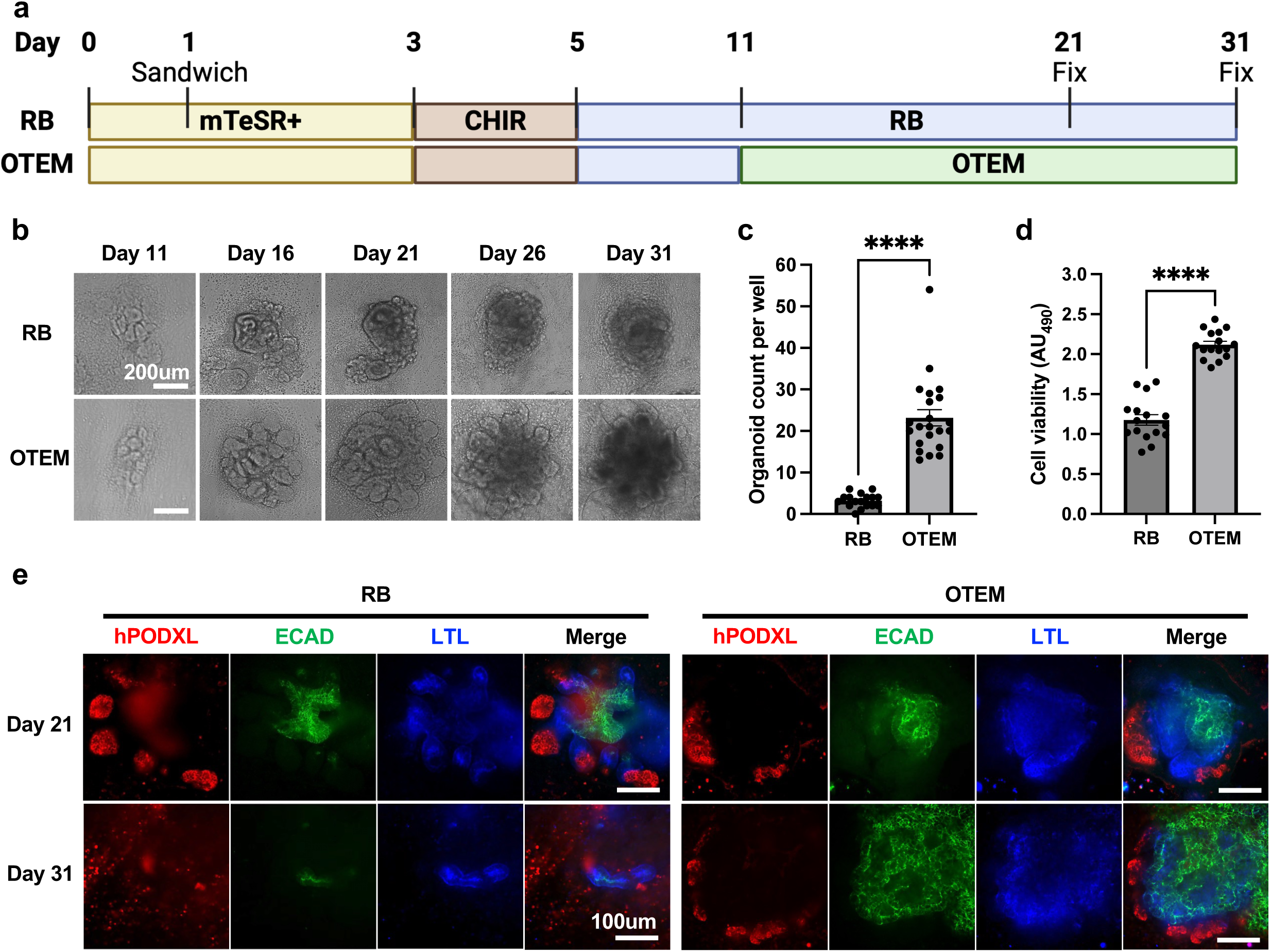
OTEM improves kidney organoid differentiation and survival. **(A)** Schematic of kidney organoid differentiation process using RB v. OTEM. **(B)** Time course brightfield images of organoids fed with OTEM starting on day 11 compared to organoids fed continuously with RB. Scale bar, 200um. **(C)** Quantification of organoid number on day 26 of differentiation in wells fed with OTEM starting on day 11 compared to wells fed continuously with RB. Mean ± s.e.m. from n >= 17 wells pooled from 5 independent experiments. Student’s t-test performed. ****, p<0.0001. **(D)** Cell viability analysis on day 26 of differentiation in wells fed with OTEM starting on day 11 compared to wells fed continuously with RB. Absorbance read at 490nm. Mean ± s.e.m. from n = 16 wells pooled from 5 independent experiments. Student’s t-test performed. ****, p<0.0001. **(E)** Comparison via confocal immunofluorescent images between organoids fed with OTEM starting on day 11 and organoids fed continuously with RB. Organoids were fixed on day 21 or day 31 and stained with ECAD (green), LTL (blue), and hPODXL (red). Scale bar, 100um. **Additional details in Supplementary Table 1 and Supplementary Figure 1.**

Compared to standard RB medium, however, OTEM substantially increased organoid yield and viability by four-fold and two-fold, respectively (**Figure 1C-D**). Whereas nephron segments of podocytes, proximal tubules, and distal tubules were well organized in both conditions at day 21, they had substantially deteriorated by day 31 in standard RB media, whereas they appeared stabilized and expanded in OTEM (**Figure 1E)**. Thus, addition of TEFs in the form of OTEM had a beneficial effect on organoid differentiation and maintenance.

In optimization experiments, we found that OTEM administration on day 11 produced the greatest increase in organoid yield and nephron segment differentiation, compared to earlier or later treatments (**Supplementary Figure 1A-C**). Increasing the concentration of TEM did not further improve organoid yield (**Supplementary Figure 1D**). Removing A83-01 from OTEM had the largest effect on organoid differentiation, significantly reducing the size of both proximal and distal tubular segments, while OTEM minus any single factor consistently produced a greater area of nephron segments than standard RB medium alone (**Supplementary Figure 1E-H**). These findings suggest that the timing of administration and the composition of OTEM contribute synergistically to the observed improvements in kidney organoid culture.

### VEFs promote kidney organoid vascularization

To promote vascularization, we tested the impact of supplementing OTEM with VEGF and ascorbic acid, a formulation we called OVA, together with gently passaging the cultures at the renal vesicle stage (**Supplementary Table 1 and Figure 2A**).^13,23,24^ Adherent organoids grown in OVA without passaging showed extensive growth of the endothelial cell network, compared to standard RB medium or OTEM, while passaged organoids further demonstrated increased invasion of organoids and podocyte clusters by the endothelial cells (**Figure 2B-G and Supplementary Figure 2A-H**). Consistently across culture conditions, endothelial cells invaded a larger proportion of podocytes than they did proximal tubules (**Supplementary Figure 2G-H**). Addition of two VEFs plus passaging thus provided a system for more advanced vascularization of kidney organoids, compared to OTEM alone.

**Figure 2.**
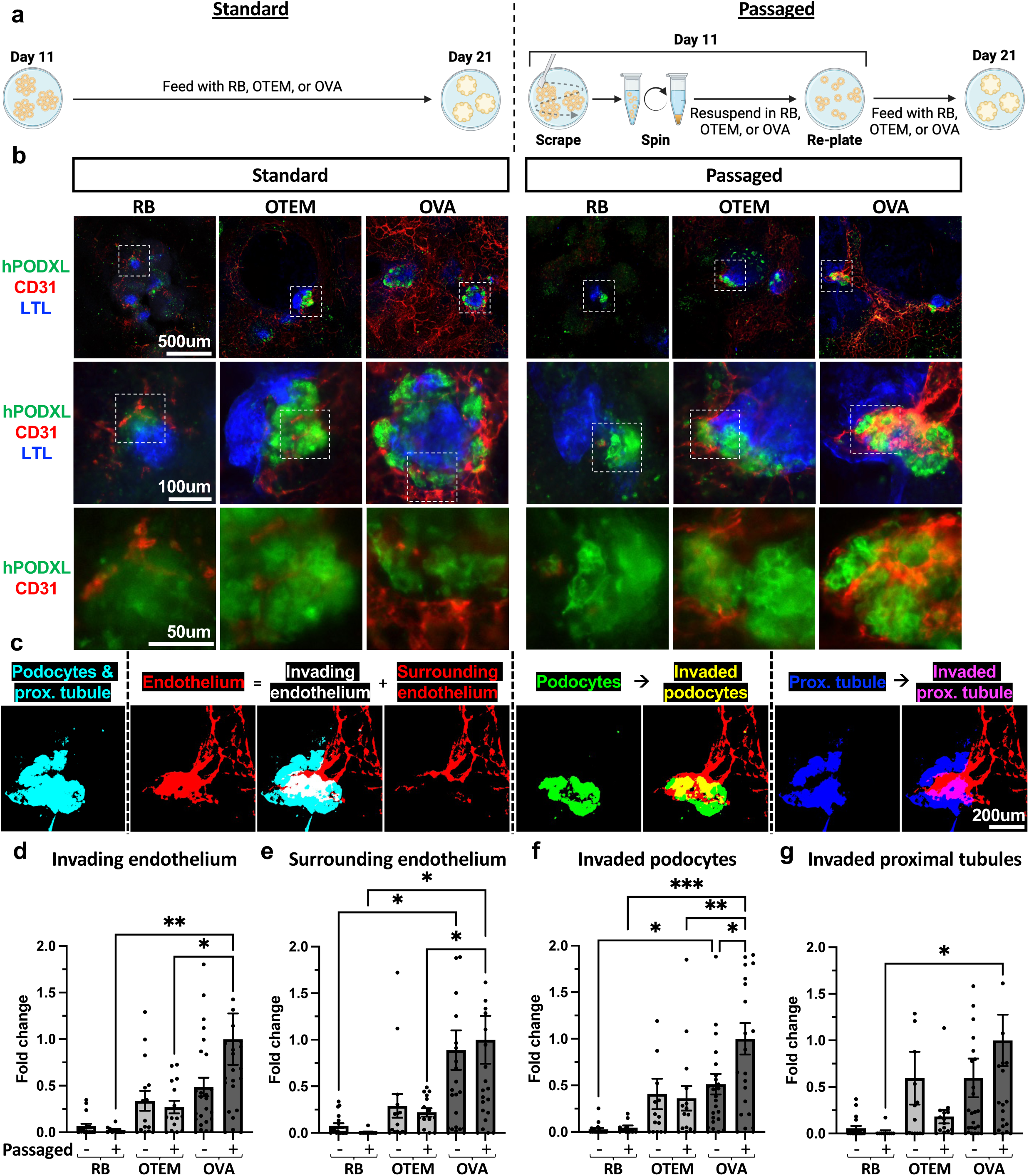
Addition of VEFs with replating promotes endothelial cell differentiation and invasion. **(A)** Schematic demonstrating the standard and passaging methods including different cell mediums used between day 11 and day 21 to induce organoid vascularization. **(B)** Macro image of organoids, including their surroundings, grown in RB, OTEM, or OVA that remained in adherent culture or were passaged on day 11. Scale bar 500um. White boxes indicate regions zoomed in on below. Scale bars 100um and 50um respectively. **(C)** Example of image quantification process using the passaged OVA organoid from panel C. The images in this panel are binary masks generated by thresholding the original image of the organoid to remove background signal and allow the area of each component of the organoid to be measured. **(D)** Invading endothelium defined as endothelial (CD31+) area overlapping with podocyte (hPODXL+) or proximal tubular (LTL+) area. **(E)** Surrounding endothelium defined as CD31+ area overlapping with neither hPODXL+ nor LTL+ area. **(F)** Invaded podocytes defined as percent of all hPODXL+ area that is overlapping with CD31+ area. **(G)** Invaded proximal tubules defined as percent of all LTL+ area that is overlapping with CD31+ area. **(D-G)** Mean ± s.e.m. from n >= 10 organoids pooled from 5 independent experiments. Outliers with fold change >2 not shown but included in analyses. Analyzed for statistical significance using a one-way ANOVA to compare passaged v. adherent within a media condition, passaged across media conditions, and adherent across media conditions. Only comparisons with p<0.05 are shown. *, p<0.05. **, p<0.01. ***, p<0.001. **Additional details in Supplementary Figure 2.**

### Transcriptomic atlas of organoid culture over 6 months

Intriguingly, a pilot study of longevity suggested that OTEM could sustain organoids for up to 6 months (**Supplementary Figure 3A**). We therefore conducted a detailed time course analysis of organoids maintained in OTEM, OVA, or standard RB media over 6 months without passaging, including characterization by single cell RNA-sequencing (scRNA-seq) and immunofluorescence at 5 different time pointss (**Figure 3A**). At day 21, organoids with well-formed podocytes (hPODXL), proximal tubules (LTL), and distal tubules (ECAD) in contiguous segments were observed in each media condition (**Figure 3B**). On days 40 and 60, organoids in OTEM and OVA maintained these nephron segments, whereas in standard RB media there was a significant degradation (**Figure 3B and Supplementary Figure 3B**). The distal tubules in both OTEM and OVA, which were originally tightly clustered around each organoid, began to elongate and span between individual organoids by day 60, a morphology that was preserved through days 120 and 180 (**Figure 3B and Supplementary Figure 3B**). At the experimental endpoint on day 180, both OTEM and OVA retained organized proximal tubule and podocyte clusters connected by long stretches of distal tubules, while the standard RB media condition had only a handful of independent tubules and islands of disorganized cells (**Figure 3B**).

**Figure 3.**
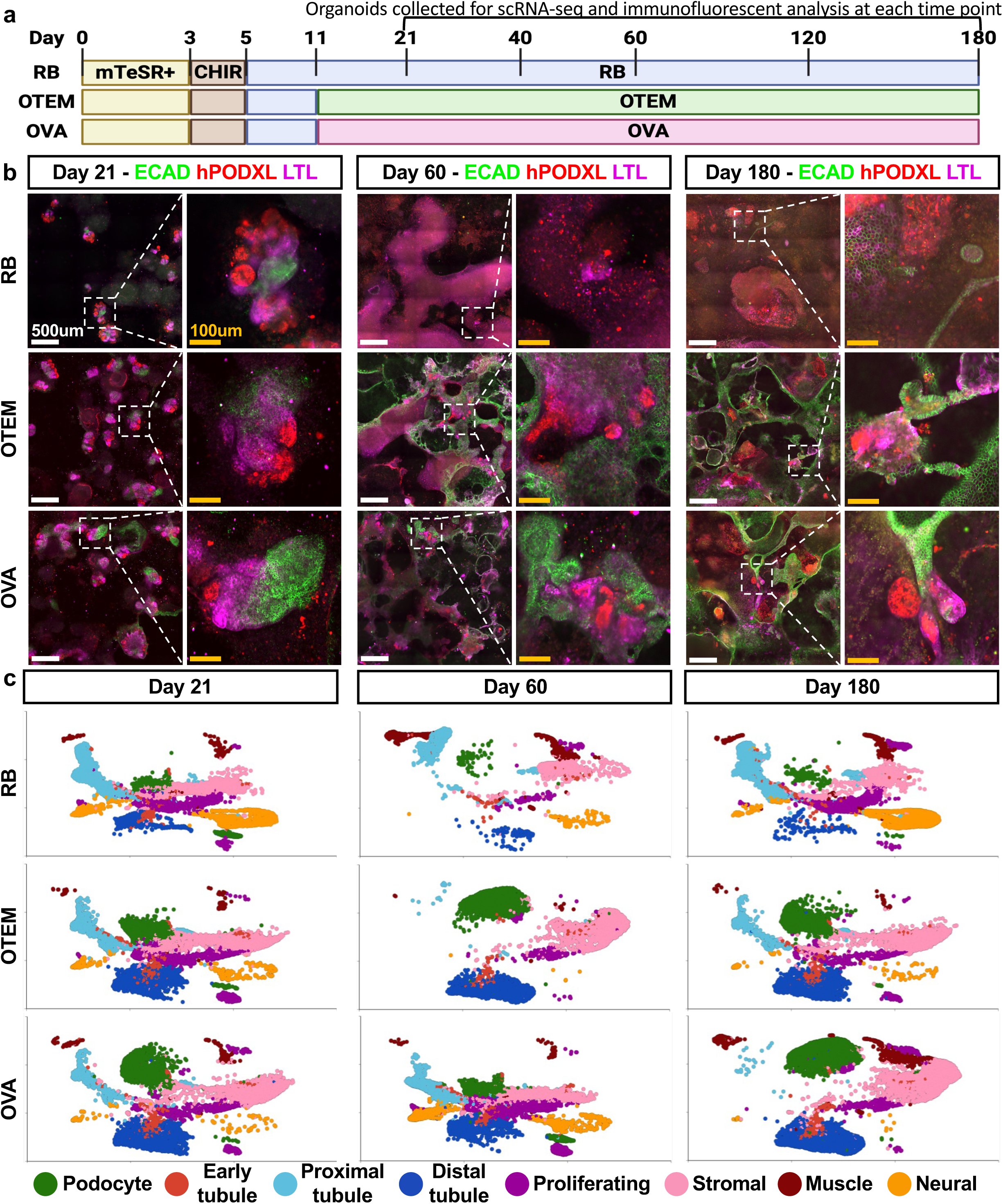
OTEM and OVA sustain kidney organoids for 6 months. **(A)** Schematic of 6-month organoid differentiation experiment. **(B)** Confocal immunofluorescent staining for classic kidney markers at days 21, 60, and 180 in all three media conditions. White box indicates the individual organoid from each larger image shown in the zoomed-in image. Scale bars are 500um and 100um, respectively. **(C)** 2D UMAPs from scRNA sequencing of each media condition across the time course. **Additional details in Supplementary Figure 3.**

To better understand how cell populations were changing over time, we generated a single-cell RNA sequencing atlas of samples from each timepoint and media condition in our longitudinal study, comprising 159,548 cells in 8 distinct clusters corresponding to podocytes, early tubules, proximal tubules, distal tubules, proliferating cells, stromal cells, muscle, or neural cells (**Figure 3C, Supplementary Figure 3C-E**). Overall, these findings demonstrated that RB, OTEM, and OVA all maintained clusters of cells with gene expression patterns characteristic of specific nephron epithelial and stromal cell populations over the six month period (**Figure 3C**). Podocytes and distal tubule segments appeared to be stabilized in OTEM and OVA, while proximal tubules - the intermediate nephron segment - appeared enriched in RB (**Figure 3C**). Thus, the deterioration of well-defined nephron segments in RB organoids observed by immunofluorescence did not reflect epithelial cell death, but rather a reduction of strong marker expression and well-defined segmented architecture, possibly merging into a dedifferentiated, proximal tubule-like state.

The overall proportion of kidney specific cell types was consistently 20-30% higher in OTEM and OVA, compared to RB, while the stromal, proliferative, and off target cell populations were more stable and lower compared to the standard RB media (**Figure 4A-D**). Although OVA increased the number of endothelial cells on day 21, this effect appeared to be diminished by day 60 (**Supplementary Figure 3E**). Endothelial cells in organoids were generally difficult to distinguish by scRNA-seq, as we have previously observed (**Supplementary Figure 3F-G**).^13,25^

**Figure 4.**
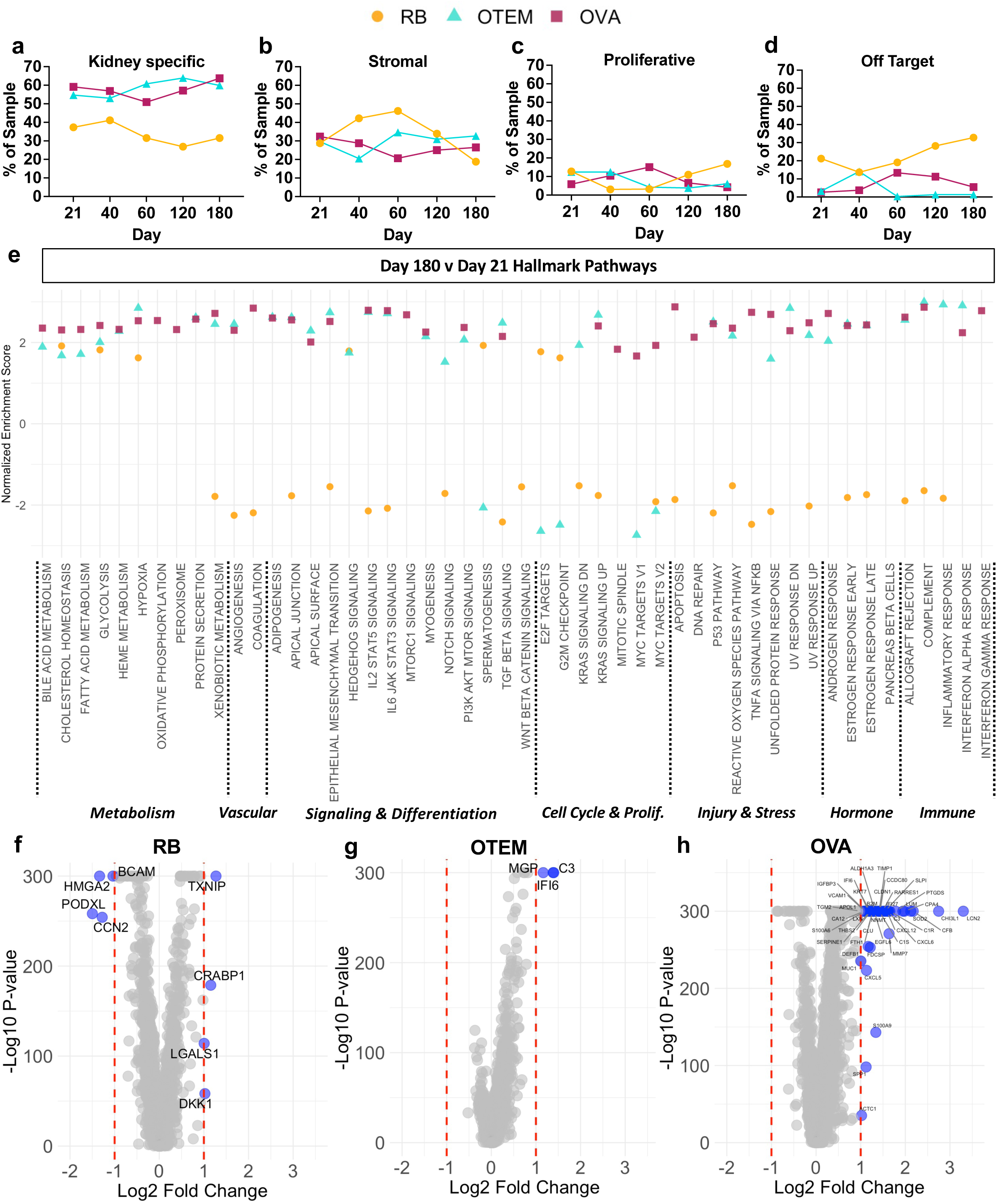
Changes in cell proportions and gene expression over time illustrate the stabilizing effect of OTEM and OVA on kidney organoids. **(A)** Quantity of kidney-specific cells as a proportion of the total sample for each media condition at each timepoint. **(B)** Size of the stromal cell cluster as a proportion of the total sample for each media condition at each timepoint. **(C)** Size of the proliferative cell cluster as a proportion of the total sample for each media condition at each timepoint. **(D)** Quantity of off target cells, muscle and neural clusters, as a proportion of the total sample for each media condition at each timepoint. **(E)** Gene set enrichment analysis (GSEA) of RB (orange), OTEM (cyan), and OVA (maroon) showing pathway enrichment at day 180 compared to day 21. Analysis was performed using the hallmark gene sets from the molecular signatures database. Data for a given media condition is only shown for a pathway if the FDR step-up score was less than or equal to 0.05. **(F)** Volcano plot of ANOVA analysis performed comparing RB at day 180 to day 21. **(G)** Volcano plot of ANOVA analysis performed comparing OTEM at day 180 to day 21. **(H)** Volcano plot of ANOVA analysis performed comparing OVA at day 180 to day 21. **Additional details in Supplementary Figure 4. (F-H)** Red dotted lines represent a Log_2_ fold change of ±1. Genes whose change in expression meets the fold change cut-offs are represented by blue dots and labelled with the gene’s name. Due to computational limits, any p-value < 1×10^−300^ is plotted as -Log_10_(1×10^−300^).

Analysis of hallmark cellular pathways, again comparing day 180 to day 21 for each media condition, revealed a broad upregulation of pathways related to cell activity, signaling, and response to stimuli in OTEM and OVA, compared to standard RB media (**Figure 4E**). Evidence of cellular senescence with prolonged culture, as assessed based upon upregulation of p53 and downregulation of E2F target genes, was increased in OTEM and OVA, compared to RB (**Figure 4E**).^26,27^ Detailed analysis of individual gene expression from day 21 to day 180 in standard RB medium revealed decreased expression of genes associated with podocytes (*PODXL*), cell adhesion (*BCAM, CCN2*), cell migration and proliferation (*HMGA2, LGALS1*); and increased expression of genes associated with Wnt/beta-catenin inhibition (*DKK1*), accumulation of reactive oxygen species (*TXNIP*), and an increasing neural population (*CRABP1*) (**Figure 4F**).^28,29^ The same analysis in OTEM revealed only three upregulated differentially expressed genes whose roles include regulating apoptosis (*IFI6*), activating the complement system (*C3*), and inhibiting tissue calcification (*MGP*) (**Figure 4G**).

Notably, there were forty-three differentially expressed genes in OVA from day 21 to day 180 associated with diverse cell activities including angiogenesis, ECM organization and cell adhesion, tumor suppression, and inflammatory/immune response (**Figure 4H and Supplementary Figure 4A**). Thus addition of just two VEFs to OTEM had a dramatic impact on gene expression patterns.

To further analyze and validate the physiological relevance of our findings, we compared the differential pathways in our d180 (aged) vs. d21 (control) OVA condition to those reported in published transcriptomic analyses of murine podocytes with renal hypertension vs. controls, or primary human proximal tubular epithelial cells undergoing induced senescence vs. controls. Five pathways were upregulated compared in the all three datasets: p53, apoptosis, inflammatory/immune pathways, TNF-alpha signaling via NFkB, and IL6-JAK-STAT3 (**Supplementary Figure 4B**). p53 activation likely reflected senescence, which along with apoptosis is associated with cell aging, while the three other pathways were suggestive of inflammation, a hallmark of chronic kidney disease. These findings validate the physiological relevance of our 180-day kidney organoid atlas for studies of renal longevity, and suggest an intrinsic ability of kidney organoids to model an immune response relevant to chronic disease, consistent with prior analyses.^30–32^

### OTEM and OVA improve organoid differentiation in diverse protocols

To investigate whether tubular and vascular enhancing factors improve outcomes in other organoid differentiation protocols and settings, we assessed these factors in a suspension-based protocol, in which derived organoids could be embedded into hydrogels mid-differentiation (**Figure 5A**). In these suspension cultures, OTEM or OVA was necessary for consistent organoid differentiation, whereas epithelial structures expressing nephron segment markers failed to form entirely in standard RB media (**Figure 5B**). In suspension, nephron-like epithelial structures consistently polarized to one side of the organoid body budding off of a large, partially hollow, stromal cell cluster (**Figure 5B-C and Supplementary Figure 5A**).

**Figure 5.**
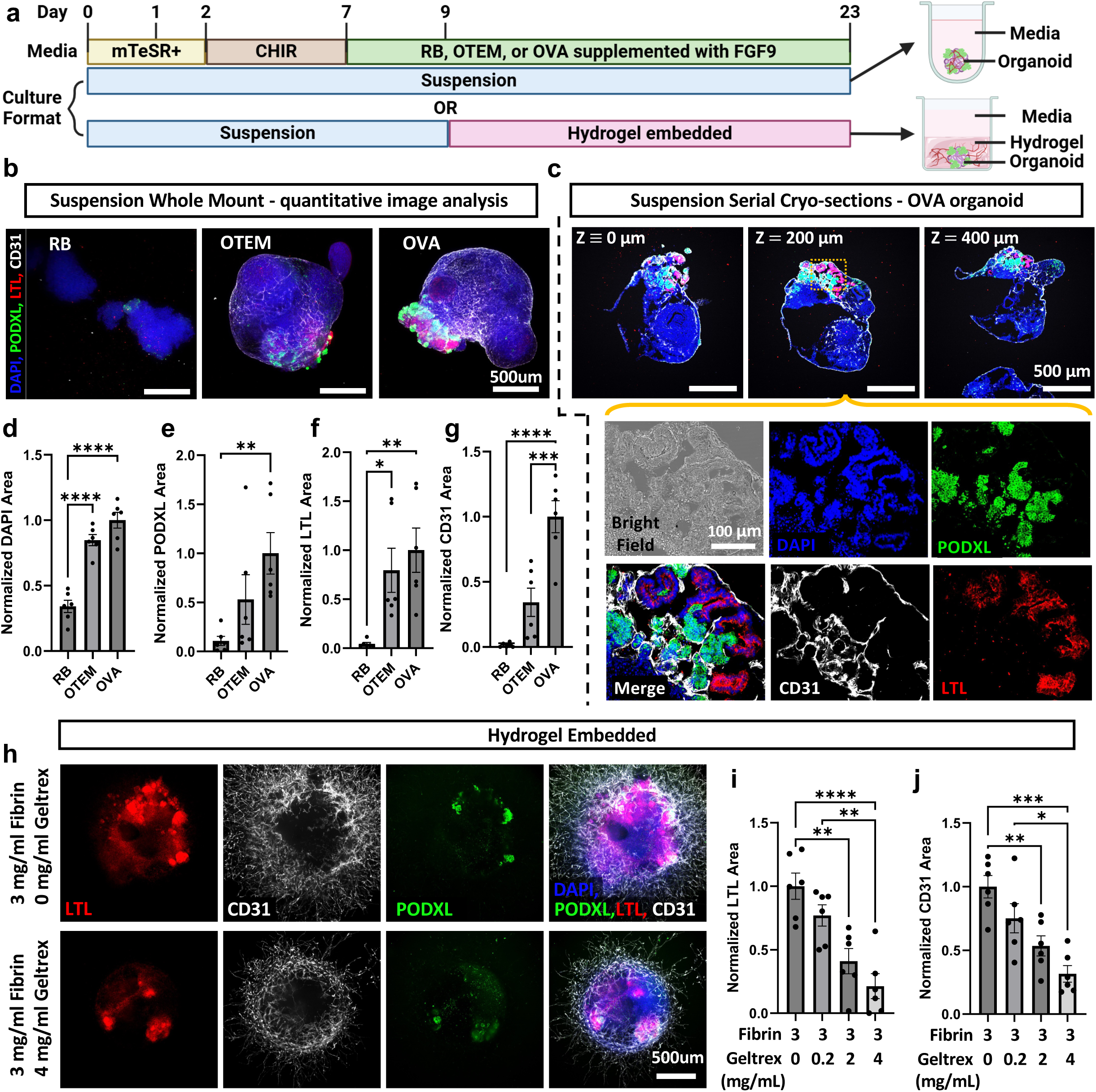
TEFs and VEFs improve organoid differentiation in suspension and facilitate embedding in hydrogels. **(A)** Schematic of adapted protocol for kidney organoid differentiation in suspension and embedding in hydrogels. **(B)** Representative maximum intensity projections of disk-spinning confocal z-stacks of whole mount suspension organoids taken at 4x magnification maintained in RB, OTEM, and OVA. Scale bars: 500 μm. **(C)** Spinning disk confocal immunofluorescent images of representative OVA-organoid serial cryo-sections at designated spacing taken at 4x. Scale bars: 500 μm. Yellow outline indicates ROI for 20x images shown below. Scale bar: 100 μm. **(D-G)** Immunofluorescence microscopy quantification of whole mount images for binary areas normalized to OVA for a given replicate showing **(D)** DAPI, **(E)** podocyte (PODXL), **(F)** proximal tubular (LTL), and **(G)** endothelial (CD31), normalized areas per organoid. Statistics as below. **(H)** Representative maximum intensity projections of disk-spinning confocal z-stacks of suspension organoids embedded in fibrin hydrogels (3 mg/ml) with varying concentrations of supplemented Geltrex. Scale bar: 500 μm. **(I, J)** Immunofluorescence microscopy quantification of binary areas normalized to 0 mg/ml Geltrex for a given replicate showing **(I)** proximal tubular (LTL), and **(J)** endothelial (CD31), areas per region of interest (4mm²) centered on an organoid. **Additional detail in Supplementary Figure 5.** For all quantification: n=6 organoids pooled between 2 independent experiments. Mean ± S.E.M. *P<0.05, **P<0.01, ***P<0.001, ****P<0.0001

Serial cryo-sectioning confirmed epithelial structure formation including the presence of ECAD+LTL- tubular regions, indicating the formation of distal tubular segments (**Supplementary Figure 5A**). CD31+ endothelial networks permeated deeply into organoids, tightly associated with podocyte tufts, and formed intimate peritubular associations (**Figure 5C**). These networks co-stained with CD146, validating endothelial identity (**Supplementary Figure 5B**). Failure of organoids to differentiate in RB was reflected quantitatively by significantly reduced DAPI area and negligible PODXL, LTL, and CD31 areas in RB compared to both OTEM and OVA (**Figure 5D-G**). Culture in OTEM resulted in organoids of comparable size and tubule formation to OVA but demonstrated inconsistent podocyte and endothelial network formation (**Figure 5D-G**), suggesting that vascularization improves podocyte differentiation.

To demonstrate the utility of OVA for bioengineering applications, we further investigated the capacity of OVA to support the differentiation of organoids cultured in fibrin-Geltrex hydrogels of varying compositions. Fibrin was chosen due to its ease of fabrication, relative affordability, and pro-angiogenic effect, and supplemented with Geltrex to temper integrin-mediated pro-inflammatory signaling associated with fibrin that might negatively affect organoid differentiation.^33,34^ Suspended cellular aggregates were embedded in hydrogels on day 9 of culture after mesoderm induction and showed immediate cellular outgrowth after embedding (**Supplementary Figure 5C**). Organoids differentiated under these conditions exhibited tubule and podocyte formation, and extensive vascular networks emanating three-dimensionally out into the surrounding hydrogels and underlying substratum (**Figure 5H and Supplementary Figure 5D**).

Despite vascular outgrowth, tight association between endothelial cells and podocytes was retained (**Supplementary Figure 5D**). Organoid size did not depend on Geltrex concentration, but there was a trend toward decreasing PODXL area, and strong dose-dependent decreases in LTL area and vascular network formation as the Geltrex concentration increased (**Figure 5H-J and Supplementary Figure 5E-F**). Overall, these results demonstrate the broad utility of our optimized media formulation in supporting kidney organoid differentiation across a variety of cell culture formats and in facilitating extensive vascular network formation into three-dimensional hydrogels.

### OVA facilitates JBM formation between organoid epithelial and endothelial cells

We further investigated whether the tight association we observed in suspension organoids between endothelial and epithelial cells included JBM formation. As a positive control for JBM immunofluorescence, we performed pan-laminin staining in healthy human kidney samples (**Supplementary Figure 6A**). Using this antibody, suspension organoids cultured in both OTEM and OVA showed strong laminin staining tightly associated with the external surfaces of epithelial structures, with increased laminin staining in OVA, likely due to improved differentiation (**Figure 6A**). Stromal regions showed comparatively faint laminin staining, despite the presence of endothelial cells, suggesting that organoid epithelial cells were primarily responsible for laminin deposition (**Figure 6A**).

**Figure 6.**
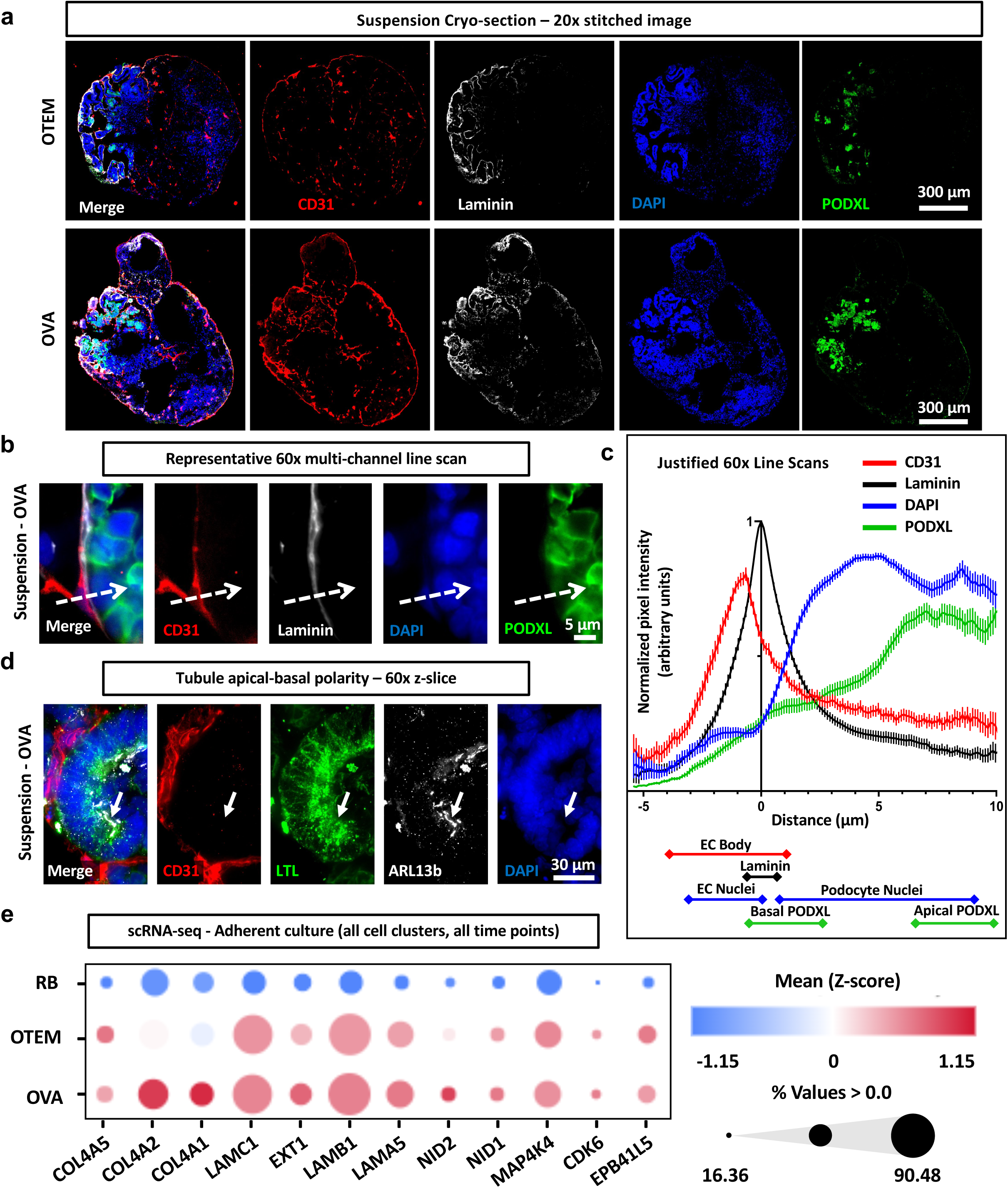
OVA facilitates JBM between organoid epithelial and endothelial cells. **(A)** Spinning disk confocal immunofluorescent images of pan-laminin expression in representative OTEM and OVA suspension organoid cryo-sections. 20x tiled image. Scale bars: 300 μm. **(B)** Representative line scan (dotted white arrow) on 60x disk-spinning confocal z-slice performed on OVA cryo-section. Scale bar: 5 μm. **(C)** Line scan quantification of background subtracted normalized fluorescence intensity across channels, justified to laminin peak intensity. Approximate demarcations of cellular structures indicated under the graph. Mean ± S.E.M. from 60 total line scans pooled across 6 organoids between 2 independent experiments. **(D)** 60x Spinning disk confocal immunofluorescent image of LTL and ARL13b expression in representative OVA suspension organoid cryo-section. White arrow indicates cilia and enriched LTL staining. Scale bar: 30 μm. **(E)** Dot plot of basement membrane gene expression from scRNA-seq of all pooled clusters and timepoints in longevity experiments comparing adherent culture in RB, OTEM and OVA. Color gradient indicates expression levels, and dot size reflects the percentage of cells expressing the gene. Gene panel based upon Maggiore et al., *J. Am. Soc. Nephrol.,* 2023. **Additional detail in Supplementary Figure 6.**

Endothelial cells formed tight associations with a thin layer of basement membrane deposited around both podocyte tufts as well as tubular structures, resembling GBMs and TBMs, respectively (**Supplementary Figure 6A**). For GBMs, line scan analysis confirmed that the laminin membrane was deposited in-between the endothelium and podocytes, with peak laminin intensity occurring between the endothelial and podocyte cell bodies, and a width on the order of single microns (**Figure 6B-C**). Expression of PODXL, a glycoprotein enriched at the apical surface of podocytes, was increased on the opposite side of the podocytes from the laminin (**Figure 6B-C**). Similarly, for TBMs, LTL staining was enriched, and primary cilia were expressed, on the surface of tubular epithelia opposite the basement membrane (**Figure 6D**). These findings demonstrated formation of GBMs and TBMs with appropriate apicobasal polarization.

We returned to our adherent organoid cultures and scRNA-seq atlas to gain additional insight into these JBMs. In replated kidney organoids cultured in OVA, we detected evidence of GBMs by immunofluorescence, although the Geltrex coating introduced some background staining, and separation between endothelial cells and laminin was difficult to discern (**Supplementary Figure 6B**). Querying our scRNA-seq atlas for a panel of genes associated with basement membrane organization revealed a robust increase in OVA compared to standard RB media, including a variety of specific collagen and laminin isoforms that contribute to GBMs and TBMs *in vivo* (**Figure 6E** and **Supplementary Figure 6C**). Collectively, our findings support the ability of kidney organoids to form JBMs *de novo* in static cultures, when provided with the appropriate microenvironment.

## DISCUSSION

### Longitudinal studies of organoids relevant to chronic disease

Deterioration over time poses a significant barrier to longitudinal studies in many different types of microphysiological systems. Typically, kidney organoids are not kept in culture for more than two months due to stromal overgrowth and structural degradation.^3–5^ The media known as TEM has recently been shown to enable serial passaging of kidney organoids derived from hPSCs for a few months.^4^ Here, we find that OTEM, a combination of TEM with our standard organoid differentiation media, maintains nephron segments *in situ* for up to six months without passaging. TEM includes RSPO3, a cofactor of Wnt/beta-catenin signaling responsible for embryonic development and tissue maintenance, and EGF, an agonist of PI3K/AKT/mTOR signaling which plays a role in maintaining stem cell pluripotency and promoting cell survival^7,35–38^. Both pathways are increased by OTEM during long-term culture, which may contribute to improved stability of the differentiated organoids.

The successful long-term culture of kidney organoids for up to six months creates new possibilities for studying chronic kidney diseases, aging, and exposures. Aging is a major risk factor in chronic kidney disease, and is associated with senesecence and inflammatory phenotypes in proximal tubular epithelial cells and podocytes^26,27^, with similarities in gene expression patterns to our aged organoids. Our atlas of longitudinal gene expression provides a window into the changes that hPSC-derived organoids *in vitro* undergo over time, an area that remains poorly understood and may be relevant to various organ lineages and disease states, such as Alzheimer’s disease for cerebral organoids^39^. In addition, the extended longevity of these organoids could enable studies of damage caused by prolonged exposures to drugs such as cisplatin, immunosuppressants, and antiretrovirals. Of note, the atlas suggests that epithelial segments may remain viable in standard RB media, but adopt a dedifferentiated state, with possible relevance for studies of kidney injury and maladaptive repair.^3,40,41^

### Self-assembly of JBMs in ‘one-pot’ format

Remarkably, addition of just two VEFs to OTEM enhances endothelial differentiation and invasion into kidney organoids, enabling JBM formation between endothelial cells and nephron epithelia. Several features are notable here. First, the JBM forms between cells with opposite apicobasal polarity, at their common basal surfaces, with no intermediate materials, and therefore constitutes a physiologically relevant basement membrane. Moreover, JBMs self-organize in static cultures, without any top-down bioengineering or flow devices. This differs from previous studies, which have used a variety of methods, including flow devices or VEGF treatments, to promote endothelial interactions with organoid epithelia, but have not clearly demonstrated a JBM with dimensions, polarity, and differentiation state that approximates the physiological condition.^13,14,42,43^ The difference here appears to be the use of both TEFs and VEFs in combination, which enables the JBMs to form.

The endothelia arise intrinsically within the organoid cultures, which simplifies the process, compared to studies wherein the endothelia are added from a separate population, which may involve genetic modification to promote its differentiation^44^. Our scRNA-seq analysis reveals that OTEM stimulates a variety of signaling pathways including mTOR, which is upstream of *HIF1,* overexpression of which has been previously shown to improve vascularization in kidney organoids.^38,45^ Epithelial cells deposit laminin as an initial step in JBM formation, which likely provides an attactive surface for vascular cells to migrate onto and coat^10^.

As the JBM is a fundamental and pervasive feature of renal tissue, which is involved in many aspects of renal function, the inability to form a JBM *in vitro* has posed a significant challenge for modeling renal physiology and disease.^9,12,46^ The improved vascular network and formation of JBM in organoids differentiated in OVA may therefore lead to functional assays *in vitro,* for instance to model drug absorption or secretion. Bioengineering approaches have been applied to mold parallel channels for endothelial cells and tubules, but these cells are physically separated by ∼100 µm of extracellular matrix, orders of magnitude larger than JBMs^16–18^. This challenge is also relevant to other organ lineages, such as the brain, where attempts have been made to reconstruct the blood-brain barrier by merging vascular and brain organoids^47^. We show that simply adding the general VEFs to the media, in combination with TEFs, is sufficient to promote JBMs to form naturally in cell culture, thus this concept is potentially widely applicable in many different types of organoids.

Although nephron segments can be maintained in long-term culture, they changed significantly over six months. In particular, the distal tubular compartment undergoes substantial expansion and outgrowth, which may reflect an interaction of this segment with the adherent substrate immediately beneath it, as well as a tendency of TEM to favor distal tubule fate over time.^7,48^ Similarly, while we have succeeded in increasing the number of endothelial cells with OVA, they fail to persist beyond the initial period of differentiation. This is consistent with the rather delicate nature of organoid endothelial cells, and demonstrates the need for additional microenvironment optimization to promote the longevity of the vasculature.^15^ It is possible that long-term exposure to VEFs is detrimental, or alternatively that the endothelial cells migrate onto the substrate and dedifferentiate, since these experiments were performed exclusively in adherent cultures. Of note, some of these features may represent processes of aging and/or inflammation during the long-term cultures. Indeed, we have recently observed that endothelial cells rapidly disappear in organoids treated even transiently with interferon gamma, an activator of JAK-STAT signaling^32^, which is a pathway we have observed to be upregulated in long-term cultures by scRNA-seq.

To date, our understanding of JBMs in these cultures is limited to structural and gene expression analyses. The vascular network growth and formation of JBM in organoids differentiated in OVA may therefore lead to *in vitro* functional assays *in vitro,* for instance to model drug absorption or secretion. Because the endothelium does not fully envelop the organoids in a confluent monolayer, but rather forms a web-like network surrounding it, there may be leakage of solutes into organoids through discontinuous regions of the endothelial network. Prolonged culture of suspension organoids, or incorporation of genetic manipulations such as *ETV2* overexpression,^44^ may promote formation of a more complete endothelial shell.

Based on our current findings, we would expect the apical surface of this endothelium to polarize outwards towards the media, while the organoid epithelium it surrounds would polarize inwards into the organoid body (apical-in). This configuration would make the media column analogous to the interior of a renal capillary. If so, transport processes may be directly observed by adding in small molecule cargoes to the surrounding media, using methodologies similar to those we have previously developed for undifferentiated iPSCs and adherent kidney organoids lacking JBMs.^11,46^ Ultimately, the goal would be to connect this vascular surface with blood vessels carrying solutes into and out of the organoids, similar to the way kidney tissues function *in vivo.* Placing OVA-differentiated organoids into microfluidic devices bearing conduits for functional blood vessels may enable such studies, resulting in a human *in vitro* system with the ability to model renal transport processes and possibly even glomerular filtration.

Much of the above remains beyond the scope of this initial study, which spans parallel advances in organoid longevity and vascularization *in vitro*. Surprisingly, limitations in both of these areas can be overcome in static cultures simply by optimizing the microenvironment. These studies, together with the associated transcriptomic longitudinal atlas in various media conditions, offer a diverse resource for long-term culture and tissue engineering applications.

## METHODS

### Stem cell culture

All work with hPSCs was conducted with approval under the auspices of the University of Washington ESCRO. Commercially available, anonymized cell lines were utilized for the study, thus the work did not constitute human subjects research. WTC11 iPSCs (UCSFi001-A), derived from a male donor, and H9 ES cells (WAe009-A), derived from a female donor, were used to differentiate kidney organoids. Both cell lines were fed 2mL of mTeSR Plus medium (100-0276, STEMCELL Technologies) every other day, supplemented with 1% penicillin/streptomycin (15140-122, Gibco) in 6-well plates (353046, Falcon) coated with 1% Geltrex (A14133-02, Gibco). Cells were incubated at 37° C and 5% CO_2_. When not being used to start a new organoid differentiation, passaging was performed using ReLeSR (100-0483, STEMCELL Technologies).

### Human kidney tissue

Cryoblocks of normal human kidney tissue were obtained from UW Pathology services. As these were completely deidentified, this did not constitute human subjects research.

### Kidney organoid differentiation from iPSCs in adherent culture

Both WTC11 and H9 stem cells were differentiated into kidney organoids using a previously published protocol for differentiation in adherent culture.^34^ Briefly, tissue culture plates were precoated with 1% GelTrex for 1 hour at 37° C. Then, stem cell colonies were dissociated using 1 mL Accutase (07922, STEMCELL Technologies) per well in 6-well plates. WTC11 cells were seeded at 100-300 cells per well in a 96-well plate (655090, Greiner Bio-One) or 1000-2000 cells per well in a 24-well plate (92024, TPP). H9 cells were seeded at 800-1200 cells per well in a 96-well plate. Stem cells were seeded in mTeSR plus medium supplemented with 10 uM Y-27632 ROCK inhibitor (1254, Tocris). 24-hours after seeding, cells were sandwiched with 1.5% GelTrex in mTeSR plus medium. Cell medium was replaced again the following day. On the evening of day 3, WTC11 spheroids were treated with 12uM CHIR 99021 (4423, Tocris) in Advanced RPMI 1640 medium (12633-012, Gibco, aRPMI) for 36 hours. H9 spheroids were treated with 10uM CHIR and 10ng/mL Noggin (120-10C, Peprotech) in aRPMI for 36 hours. Culture medium was then replaced with RB: aRPMI supplemented with 1% penicillin/streptomycin, 1X Glutamax (35050-06, Gibco), and 1X B27 supplement (17504044, Gibco) and replaced every other day for the duration of each experiment.

### Organoid Tubuloid Enhancing Media (OTEM) and OVA treatment

Kidney organoids were treated with OTEM between days 5 and 16. OTEM was prepared by supplementing standard differentiation media with tubuloid expansion media at a 1:5 ratio (**Supplementary Table 1**). Tubuloid expansion media, as previously described, is composed of: 94% Advanced DMEM/F12 (12634-010, Gibco), 1% penicillin/streptomycin, 1% HEPES buffer (83264, Sigma-Aldrich), 1% Glutamax, 0.1mg/mL primocin (ant-pm-1, InvivoGen), 1.6% B27 supplement, 1% R-Spondin 3 conditioned medium (SCM105, Millipore Sigma), 50 ng/mL EGF (AF-100-15-500UG, Peprotech), 100 ng/mL FGF-10 (345-FG-025, R&D Systems), 1mM N-acetyl-L-cystine (A9165-25G, Sigma-Aldrich), 5 uM A83-01 (2939, Tocris), and 10mM Y-27632 ROCK inhibitor. ^4,7,22^ Different ratios of tubuloid expansion media to standard differentiation media were generated by diluting the same tubuloids expansion media in varying amounts of RB until the desired concentration was reached. Organoids continued to be fed every other day once OTEM treatment began.

OVA consisted of OTEM was supplemented with 1.05 uL/mL VEGF (LS-1016, Lifeline Cell Technology) and 1.05 uL/mL ascorbic acid (LS-1005, Lifeline Cell Technology) (**Supplementary Table 1**). Treatment with this media began on day 11 of differentiation and media was changed every other day through the end of an experiment.

### Kidney organoid passaging

Passaging kidney organoids was always performed on day 11 of differentiation, exclusively in 24-well plates. First, culture media was aspirated until only 500uL remained in each well to be passaged. Then, a mini cell scraper (MCS-200, ABI Scientific) was used to mechanically lift the differentiating organoids from the well, scraping in a 5×5 grid pattern and a circular sweep around the edges of the well. The cell suspension was aspirated using a p1000 pipet and transferred to a 1.5mL microcentrifuge tube (229442, CellTreat). The contents of each well were transferred into a separate tube. These were centrifuged for 4 minutes at 1100 rpm, the supernatant was aspirated, and the differentiating organoids were resuspended in either RB, OTEM, or OVA before being re-plated into a new 24-well plate that was pre-coated with 1% GelTrex. Organoids were fed every other day after passaging through day 21.

### Suspension Organoid Differentiation

The suspension-based protocol was adapted from an established protocol using media from our standard differentiation protocol. Briefly, WTC11 iPSCs were maintained as described for our standard differentiation protocol and grown to ≥ 75% confluence prior to starting the kidney differentiation protocol. On day 0, cells were rinsed with PBS then incubated in 1 ml ACCUTASE (7922, STEMCELL Technologies) in the incubator until cell detachment, pelleted at 500G, and resuspended at a concentration of 100,000 cells/mL in mTeSR+ supplemented with 10 μM Y-27632 dihydrochloride (1254, Tocris Bioscience). 200 μL of cell suspension was added per well of ultra-low attachment round bottom 96-well microplate (CLS7007, Corning). On day 1, medium in each well was replaced with mTeSR+ without Y-27632 dihydrochloride. On day 2, medium was replaced with advanced RPMI 1640 medium (aRPMI) (12633020, Thermo Fisher) supplemented with 9μM CHIR 99021 (4423, Tocris Bioscience), 2 mM GlutaMAX (35050061, Thermo Fisher), and 100 U/mL Pen-Strep (15-140-12, Thermo Fisher). This media was replaced on days 3 and 5. On day 7, medium was replaced with RB, OTEM, or OVA. On day 9, organoids were either embedded in hydrogels or continued in suspension culture in RB, OTEM, or OVA supplemented with 10 ng/mL recombinant human FGF9 (273-F9-025/CF, R&D Systems). Media was exchanged every 2-3 days for the remainder of culture and organoids were fixed on day 23. Cells were maintained at 37 °C and 5% CO2 in a medium volume of 200 μL/well throughout the differentiation protocol.

### Hydrogel Formulation and Suspension Organoid Embedding

Fibrin/Geltrex composite hydrogels were formulated with constant final fibrin concentration (3 mg/mL clottable protein), varying final Geltrex concentration (0 – 4 mg/mL), and aprotinin (7.5 μM) to prevent cellular- and Geltrex-mediated fibrin degradation. Lyophilized fibrinogen (F8630, Sigma-Aldrich) was dissolved in DMEM/F-12, vortexed extensively, degassed, and sterile filtered with a 0.2 micron syringe filter (09-719C, Fisherbrand). Polymer precursor solution containing Fibrinogen, aprotinin, and Geltrex was kept on ice until hydrogel casting and polymerized with 1U/mL final concentration of thrombin (T7513, Sigma-Aldrich). Immediately prior to hydrogel casting, organoid media was aspirated and replaced with thrombin diluted in DMEM in the U-bottom plate. Hydrogels were cast into 96-well TC treated plates (655090, Greiner Bio-One) by gently mixing polymer precursor solution with thrombin and the organoid in the U-bottom plate, then transferring with a wide-orifice 200 μL pipette tip. All hydrogels were 80 μL, cast with a single organoid per gel and allowed to polymerize at 37 °C and 5% CO2 for 45 minutes before addition of culture media. Organoids were maintained in a medium volume of 200 μL/well supplemented with aprotinin (7.5 μM) throughout the differentiation protocol.

### Quantification of kidney organoid differentiation yield

The number of organoids per well was determined by taking live brightfield whole well images at 4x magnification using an Olympus IX83 microscope equipped with a disk spinning unit (DSU, Evident) and cellSense software. The number of organoids in each well were then counted manually.

### Viability assay

Cell viability was quantified using the CellTiter 96 AQ_ueous_ One Solution Cell Proliferation Assay (G3582, Promega). The assay was performed following the manufacturer’s protocol. Briefly, for one well of a 96-well plate, 20uL of the CellTiter 96 AQ_ueous_ One Solution was mixed with 100uL of DMEM/F12 medium (11320082, Thermo Fisher). Then, the culture medium was removed from wells and replaced with the prepared reagent mixture. The plate was incubated at 37° C for 1 hour, then absorbance readings were taken at 492nm on a Perkin Elmer Envision 2104 Multilabel reader.

### Immunofluorescent staining and imaging of adherent organoids

Adherent kidney organoids were fixed in 4% paraformaldehyde (15710, Electron Microscopy Sciences) in PBS (10010-023, Gibco) for 15 minutes at room temperature. They were then washed 3 times in PBS for 5 minutes each and stored at 4° C until the next step. Samples were blocked for 1 hour at room temperature with blocking buffer: 94.7% PBS, 5% donkey serum (S30-100ML, Millipore Sigma), and 0.3% Triton X-100 (1086431000, Millipore Sigma). Blocking buffer was centrifuged for 2 minutes at 3,000x*g* before use. Primary antibodies were diluted in antibody dilution buffer (ADB): 96.7% PBS, 3% bovine serum albumin (BP9703-100, Fisher), and 0.3% Triton X-100. The mixture was centrifuged for 2 minutes at 3,000x*g,* added to the organoid samples, and incubated overnight at 4° C. Samples were washed 3 times with PBS for 5 minutes each. Secondary antibodies and 4′,6-diamidino-2-phenylindole (62248, Thermo Fisher, 1:200, DAPI) were then diluted in ADB and the mixture was centrifuged for 2 minutes at 3,000x*g*, added to the organoid samples, protected from light, and incubated overnight at room temperature. Samples were washed 3 times with PBS for 5 minutes each, protected from light to prevent photobleaching, and stored in PBS at 4° C until being imaged. Confocal images were acquired using an Olympus IX83 microscope equipped with a disk spinning unit (DSU, Evident) and cellSense software.

*Primary antibodies:* anti-E cadherin (AB11512, Abcam, 1:300, ECAD), anti-podocalyxin (AF1658, R&D systems, 1:500, hPODXL), anti-LTL (B-1325, Vector Labs, 1:500, LTL), anti-laminin (AB11575, Abcam, 1:300), anti-ARL13b (17711-1-AP, Proteintech, 1:300), anti-CD146 (AB75769, Abcam, 1:300)

*Secondary antibodies:* anti-CD31-PE (REA1028, Miltenyi Biotec, 1:300), AlexaFluor 488 anti-goat (A11055, Invitrogen, 1:500), AlexaFluor 488 anti-rat (A21208, Invitrogen, 1:500), AlexaFluor 555 anti-goat (A21432, Invitrogen, 1:500), AlexaFluor 647 streptavidin (S32357, Invitrogen, 1:500)

### Image analysis

Any images that were directly compared to one another were acquired using the same staining and imaging conditions to avoid introducing bias. After image acquisition, in-house IJ1 Macros were used to analyze the images and generate raw data. Microsoft Excel was used to process the data and GraphPAD Prism was used to visualize the data and perform statistical analyses. The raw ImageJ script, written as an IJ1 Macro, is available as a supplementary methods section with accompanying notes on necessary user edits for adaptation.

### Full well ECAD/LTL/hPODXL area calculations

A full-well 4x confocal image was taken, then split into individual channels. A threshold was applied to each channel to remove background signal and highlight the regions of interest. Each channel’s threshold values were kept constant across the full set of images being quantified for a given experiment. The thresholded area was then calculated using an IJ1 Macro.

### Endothelial, podocyte, and proximal tubule measurements in individual organoids

10x z-stack images were taken, then a FIJI script was used to extract individual slices in 10% increments moving from the bottom to the top of the organoid. The area of each channel individually was quantified as described above. Again, each channel’s threshold values were kept constant across the full set of images being quantified for a given experiment. In addition, for these calculations FIJI’s image math calculator was used to quantify the overlap between endothelial cell area (CD31+) and the other organoid segments (hPODXL+ and LTL+). Quantification for **Figure 2D-G and Supplementary Figure 2A-H** was performed on a single image slice taken midway through the height of the organoid.

### Single-cell RNA sequencing

Kidney organoids differentiated in 96-well plates were treated with RB, OTEM, or OVA starting on day 11 until the cells were collected for sequencing. 20 wells of each media condition at each time point were dissociated into single cell suspensions for sequencing purposes. The time points collected were day 21, day 40, day 60, day 120, and day 180. Cells were dissociated using 200 uL of trypLE select 1x (12563-011, Gibco) per well then incubated for 20 minutes at 37° C. The dissociation was mechanically assisted by fluxing the trypLE 5-10 times in each well every 5 minutes during the incubation period. After the cells fully dissociated, the 20 wells of each condition were combined in 10mL of RB with 1% BSA to quench the trypLE and generate 3 distinct samples per time point. Samples were then centrifuged for 4 minutes at 1100 rpm, the supernatant was aspirated, and the cells were resuspended in 1 mL RB. Samples were filtered using a 40uM cell strainer (25-375, Genesee Scientific) before centrifuging again for 4 minutes at 1100 rpm, aspirating the supernatant, and resuspending in 1mL Fixation Buffer from the Chromium Next GEM Single Cell Fixed RNA Sample Prep Kit (1000414, 10X Genomics). After a 24hr incubation at 4° C, the manufacturer’s protocol for the sample prep kit was followed to prepare samples for long-term storage at -80° C.

Once samples for all time points had been collected, fixed, and stored at -80° C, the scRNA sequencing process began. Single cell droplet libraries were generated in the 10X Genomics Chromium controller according to the manufacturer’s instructions in the Chromium Fixed RNA kit, human transcriptome probe set, 4rxns x 4BC (1000475, 10X Genomics). Additional components used for library preparation included the Chromium Next GEM chip Q single cell kit, 16rxn (1000422, 10X Genomics), SPRIselect beads (B23317, Beckman Coulter), and Dual Index Plate TS Set A (1000251, 10X Genomics). Final libraries were pooled at 2nM and sequenced on an Illumina NextSeq2000 using a NextSeqTM 2000 P4 XLEAP- SBSTM Reagent Kit (20100994, Illumina, 100 Cycles). Sequencing parameters were selected according to the Chromium Fixed Single Cell specifications.

### Bioinformatic analysis of sequencing data

Sequencing base calls were converted to FASTQ files demultiplexed by sample indexes using Illumina’s BaseSpace DRAGEN Analysis Application v1.2.1. FASTQ files were uploaded to 10x Genomics’ Cloud Analysis platform and processed using the Cell Ranger Count Pipeline v7.0.1. Data are available at the National Center for Biotechnology Information’s Gene Expression Omnibus (GEO), accession number GSE286583.

Resultant feature/cell matrix (filtered) files were imported to Partek Flow for quality filtering (cells with UMI < 26,000, genes < 8,055, and ribosomal counts < 45% were kept). Counts were normalized using TMM (Trimmed mean of M-values) for DESeq2 differential expression analysis with default parameters. For UMAP representation and cluster analysis data was normalized using SCTransform and principal component analysis (PCA) was performed on the output SC scaled data followed by UMAP (Uniform Manifold Approximation and Projection) dimension reduction or graphed-based clustering under default settings.

UMAP representation and cluster analysis was performed on the full dataset first, before being split by timepoint or media condition for differential gene expression analyses.

Volcano plot data was generated by performing an ANOVA on each media separately using day 180 as the numerator and day 21 as the denominator for the analysis to output data regarding individual genes. Hallmark pathway enrichment scores were generated by performing a gene set enrichment analysis (GSEA), again on each media separately using day 180 as the numerator and day 21 as the denominator, using the Human_MSigdb_December_01_2022_symbol.gmt version of the MSigdb database (www.gsea-msigdb.org). The FDR step-up cut-off was set to be <0.05. The visual representations of both volcano plots and differentially expressed hallmark pathways were generated using R Studio and the scripts are available as a supplementary methods section with accompanying notes on necessary user edits for adaptation.

Comparisons of Hallmark pathways and GSEA analyses between studies were performed through manual extraction and measurements of pathway data rendered in graphical format from each study and their associated normalized enrichment scores. Graphs were generated using GraphPad Prism 10.4.1.

### Suspension Organoid Cryo-sectioning

Organoids for cryosectioning were fixed in 4% PFA, equilibrated over night in 30% sucrose in PBS, and then embedded in OCT (Tissue-Plus, Thermo Fisher Scientific). Embedded organoids were cryosectioned at 20 μm thickness and mounted on superfrost slides (Fisherbrand), being careful to track section number and maintain orientation, then stored at -80°C until staining.

### Suspension Organoid Semi-Automated Quantitative Immunofluorescence Microscopy

Suspension organoids were fixed in 1:1 ratio of culture media and 8% PFA (4% working solution) for 15 minutes at room temperature. Suspension organoids embedded in hydrogels were rinsed for 20 minutes in PBS then fixed with 4% PFA for 15 minutes at room temperature. After fixing, samples were stained as described above where details on antibodies used including sourcing and dilutions can also be found.

For quantitative image analyses, images were acquired using an Olympus IX83 microscope equipped with a disk spinning unit (DSU, Evident) and cellSense software. To ensure the full thickness of the organoid and surrounding hydrogel was imaged, 600 μm thick confocal z-stacks (50 μm/slice) were acquired using the DSU, imaged at 4x. Representative confocal z-slices of individual epithelial tufts were taken at 20x or 60x.

Semi-automated image analysis was performed in ImageJ for quantification of relative organoid areas (DAPI), tubular areas (LTL), podocyte areas (PODXL), and endothelial network areas (CD31). For all analyses, 4x Z-stacks were collapsed and quantification was performed on their maximum intensity projections. Accurate quantification of fluorescent images depends on uniform staining and imaging across conditions, as such, all conditions within a given replicate were identically stained and all imaging for a set was performed within 1-2 days to minimize the chance of signal fading. Relative areas for each marker were determined by uniformly thresholding all images within a replicate to generate binaries, binary areas per organoid were then normalized for each replicate. The raw ImageJ script, written as an IJ1 Macro, is available as a supplementary methods section with accompanying notes on necessary user edits for adaptation.

Line scan analysis was performed on 60x confocal z-slices. A line scan was drawn in regions that contained clear endothelial cell bodies sharing a joint laminin basement membrane with a podocyte cluster. Line scans were drawn in the direction from endothelial cells towards podocytes. The line was drawn perpendicular to the membrane and through the center of the podocyte nuclei. If an endothelial nucleus was discernable, the line was also drawn through the center of the EC nuclei. Each individual line scan was background subtracted (minimum fluorescent intensity value), normalized, and justified; the maximum laminin intensity was defined as the zero point on the x-axis for a given multichannel scan.

### Quantification and statistical analysis

Statistical analyses were performed using GraphPad Prism software. The details regarding number of replicates, graphical data representation, and specific statistical tests can be found in each corresponding figure legend.

## Supporting information

Blackburnetal_Supplement

## DATA AVAILABILITY

The main data supporting the results in this study are available within the paper and its Supplementary Information. Complete scRNA-seq data are available from a public repository. The raw and analysed datasets generated during the study are too large and complex to be publicly shared (numerous cell lines, replicates, images, and experiments, maintained and analysed in specialized file formats and with unique identifiers), yet they are available for research purposes from the corresponding author on reasonable request. Source data are included for analyses of scRNA-seq datasets (differential gene expression and pathways).

## CODE AVAILABILITY

ImageJ scripts are attached to this paper as Supplementary Information. scRNA-seq analysis code is available on request from the corresponding author.

## ACKNOWLEDGMENTS

We thank Anton Kary and Jessica Ayers for technical contributions to preliminary studies, Kelly Smith for donation of healthy human kidney pathology samples, and Edward Kelly, Catherine Yeung, Jonathan Himmelfarb, Stuart Shankland, Hongxia Fu, Ying Zheng, Seiji Kishi, and members of the Freedman lab for helpful discussions. Some illustrations were created with the assistance of BioRender.com under a paid license. Studies were supported by NASA contract 80ARC023CA001 (B.S.F.), NIH awards UC2DK126006 (B.S.F.), UH3TR003288 (B.S.F.), U2CTR004867 (B.S.F.), a Washington Research Foundation Technology Development Award (B.S.F.), and an Institute for Stem Cell and Regenerative Medicine Fellows Award (S.M.B.).

## AUTHOR CONTRIBUTIONS

B.S.F. conceived the project. S.M.B., B.A.J., A.S., M.C.R., and B.S.F. designed the experiments. S.M.B., B.A.J., and A.S. performed the experiments. S.M.B., B.A.J., A.S., M.C.R., and B.S.F. analyzed the data. S.M.B., B.A.J., and B.S.F. wrote the paper with input from all authors.

## REFERENCES

1. Li, M. & Izpisua Belmonte, J. C. Organoids - preclinical models of human disease. N. Engl. J. Med. 380, 569–579 (2019).

2. Digby, J. L. M., Vanichapol, T., Przepiorski, A., Davidson, A. J. & Sander, V. Evaluation of cisplatin-induced injury in human kidney organoids. Am. J. Physiol. Renal Physiol. 318, F971–F978 (2020).

3. Gupta, N., Matsumoto, T., Hiratsuka, K., Garcia Saiz, E., Galichon, P., Miyoshi, T., Susa, K., Tatsumoto, N., Yamashita, M. & Morizane, R. Modeling injury and repair in kidney organoids reveals that homologous recombination governs tubular intrinsic repair. Sci. Transl. Med. 14, eabj4772 (2022).

4. Yousef Yengej, F. A., Jansen, J., Ammerlaan, C. M. E., Dilmen, E., Pou Casellas, C., Masereeuw, R., Hoenderop, J. G., Smeets, B., Rookmaaker, M. B., Verhaar, M. C. & Clevers, H. Tubuloid culture enables long-term expansion of functional human kidney tubule epithelium from iPSC-derived organoids. Proc. Natl. Acad. Sci. U. S. A. 120, (2023).

5. Subramanian, A., Sidhom, E.-H., Emani, M., Vernon, K., Sahakian, N., Zhou, Y., Kost-Alimova, M., Slyper, M., Waldman, J., Dionne, D., Nguyen, L. T., Weins, A., Marshall, J. L., Rosenblatt-Rosen, O., Regev, A. & Greka, A. Single cell census of human kidney organoids shows reproducibility and diminished off-target cells after transplantation. Nat. Commun. 10, 5462 (2019).

6. Sato, T., Vries, R. G., Snippert, H. J., van de Wetering, M., Barker, N., Stange, D. E., van Es, J. H., Abo, A., Kujala, P., Peters, P. J. & Clevers, H. Single Lgr5 stem cells build crypt-villus structures in vitro without a mesenchymal niche. Nature 459, 262–265 (2009).

7. Schutgens, F., Rookmaaker, M. B., Margaritis, T., Rios, A., Ammerlaan, C., Jansen, J., Gijzen, L., Vormann, M., Vonk, A., Viveen, M., Yengej, F. Y., Derakhshan, S., de Winter-de Groot, K. M., Artegiani, B., van Boxtel, R., Cuppen, E., Hendrickx, A. P. A., van den Heuvel-Eibrink, M. M., Heitzer, E., Lanz, H., Beekman, J., Murk, J.-L., Masereeuw, R., Holstege, F., Drost, J., Verhaar, M. C. & Clevers, H. Tubuloids derived from human adult kidney and urine for personalized disease modeling. Nat. Biotechnol. 37, 303–313 (2019).

8. Hohenstein, B. & Hugo, C. Peritubular capillaries: an important piece of the puzzle. Kidney Int. 91, 9–11 (2017).

9. Morais, M. R. P. T., Tian, P., Lawless, C., Murtuza-Baker, S., Hopkinson, L., Woods, S., Mironov, A., Long, D. A., Gale, D. P., Zorn, T. M. T., Kimber, S. J., Zent, R. & Lennon, R. Kidney organoids recapitulate human basement membrane assembly in health and disease. Elife 11, (2022).

10. Abrahamson, D. R., St John, P. L., Stroganova, L., Zelenchuk, A. & Steenhard, B. M. Laminin and type IV collagen isoform substitutions occur in temporally and spatially distinct patterns in developing kidney glomerular basement membranes. J. Histochem. Cytochem. 61, 706–718 (2013).

11. Freedman, B. S., Brooks, C. R., Lam, A. Q., Fu, H., Morizane, R., Agrawal, V., Saad, A. F., Li, M. K., Hughes, M. R., Werff, R. V., Peters, D. T., Lu, J., Baccei, A., Siedlecki, A. M., Valerius, M. T., Musunuru, K., McNagny, K. M., Steinman, T. I., Zhou, J., Lerou, P. H. & Bonventre, J. V. Modelling kidney disease with CRISPR-mutant kidney organoids derived from human pluripotent epiblast spheroids. Nat. Commun. 6, 8715 (2015).

12. Takasato, M., Er, P. X., Chiu, H. S., Maier, B., Baillie, G. J., Ferguson, C., Parton, R. G., Wolvetang, E. J., Roost, M. S., Chuva de Sousa Lopes, S. M. & Little, M. H. Kidney organoids from human iPS cells contain multiple lineages and model human nephrogenesis. Nature 526, 564–568 (2015).

13. Czerniecki, S. M., Cruz, N. M., Harder, J. L., Menon, R., Annis, J., Otto, E. A., Gulieva, R. E., Islas, L. V., Kim, Y. K., Tran, L. M., Martins, T. J., Pippin, J. W., Fu, H., Kretzler, M., Shankland, S. J., Himmelfarb, J., Moon, R. T., Paragas, N. & Freedman, B. S. High-Throughput Screening Enhances Kidney Organoid Differentiation from Human Pluripotent Stem Cells and Enables Automated Multidimensional Phenotyping. Cell Stem Cell 22, 929–940.e4 (2018).

14. Homan, K. A., Gupta, N., Kroll, K. T., Kolesky, D. B., Skylar-Scott, M., Miyoshi, T., Mau, D., Valerius, M. T., Ferrante, T., Bonventre, J. V., Lewis, J. A. & Morizane, R. Flow-enhanced vascularization and maturation of kidney organoids in vitro. Nat. Methods 16, 255–262 (2019).

15. Ryan, A. R., England, A. R., Chaney, C. P., Cowdin, M. A., Hiltabidle, M., Daniel, E., Gupta, A. K., Oxburgh, L., Carroll, T. J. & Cleaver, O. Vascular deficiencies in renal organoids and ex vivo kidney organogenesis. Dev. Biol. 477, 98–116 (2021).

16. Chapron, A., Chapron, B. D., Hailey, D. W., Chang, S.-Y., Imaoka, T., Thummel, K. E., Kelly, E., Himmelfarb, J., Shen, D. & Yeung, C. K. An Improved Vascularized, Dual-Channel Microphysiological System Facilitates Modeling of Proximal Tubular Solute Secretion. ACS Pharmacol Transl Sci 3, 496–508 (2020).

17. Lin, N. Y. C., Homan, K. A., Robinson, S. S., Kolesky, D. B., Duarte, N., Moisan, A. & Lewis, J. A. Renal reabsorption in 3D vascularized proximal tubule models. Proc. Natl. Acad. Sci. U. S. A. 116, 5399–5404 (2019).

18. Petrosyan, A., Cravedi, P., Villani, V., Angeletti, A., Manrique, J., Renieri, A., De Filippo, R. E., Perin, L. & Da Sacco, S. A glomerulus-on-a-chip to recapitulate the human glomerular filtration barrier. Nat. Commun. 10, 3656 (2019).

19. Sharmin, S., Taguchi, A., Kaku, Y., Yoshimura, Y., Ohmori, T., Sakuma, T., Mukoyama, M., Yamamoto, T., Kurihara, H. & Nishinakamura, R. Human Induced Pluripotent Stem Cell-Derived Podocytes Mature into Vascularized Glomeruli upon Experimental Transplantation. J. Am. Soc. Nephrol. 27, 1778–1791 (2016).

20. van den Berg, C. W., Ritsma, L., Avramut, M. C., Wiersma, L. E., van den Berg, B. M., Leuning, D. G., Lievers, E., Koning, M., Vanslambrouck, J. M., Koster, A. J., Howden, S. E., Takasato, M., Little, M. H. & Rabelink, T. J. Renal Subcapsular Transplantation of PSC-Derived Kidney Organoids Induces Neo-vasculogenesis and Significant Glomerular and Tubular Maturation In Vivo. Stem Cell Reports 10, 751–765 (2018).

21. Freedman, B. S. & Dekel, B. Engraftment of Kidney Organoids In Vivo. Curr Transplant Rep 10, 29–39 (2023).

22. Gijzen, L., Yousef Yengej, F. A., Schutgens, F., Vormann, M. K., Ammerlaan, C. M. E., Nicolas, A., Kurek, D., Vulto, P., Rookmaaker, M. B., Lanz, H. L., Verhaar, M. C. & Clevers, H. Culture and analysis of kidney tubuloids and perfused tubuloid cells-on-a-chip. Nature Protocols 16, (2021).

23. May, J. M. & Harrison, F. E. Role of Vitamin C in the Function of the Vascular Endothelium. Antioxid. Redox Signal. 19, 2068–2083 (2013).

24. Bejoy, J., Farry, J. M., Qian, E. S., Dearing, C. H., Ware, L. B., Bastarache, J. A. & Woodard, L. E. Ascorbate protects human kidney organoids from damage induced by cell-free hemoglobin. Dis. Model. Mech. 16, (2023).

25. Harder, J. L., Menon, R., Otto, E. A., Zhou, J., Eddy, S., Wys, N. L., O’Connor, C., Luo, J., Nair, V., Cebrian, C., Spence, J. R., Bitzer, M., Troyanskaya, O. G., Hodgin, J. B., Wiggins, R. C., Freedman, B. S., Kretzler, M., European Renal cDNA Bank (ERCB) & Nephrotic Syndrome Study Network (NEPTUNE). Organoid single cell profiling identifies a transcriptional signature of glomerular disease. JCI Insight 4, (2019).

26. McKinzie, S. R., Kaverina, N., Schweickart, R. A., Chaney, C. P., Eng, D. G., Pereira, B. M. V., Kestenbaum, B., Pippin, J. W., Wessely, O. & Shankland, S. J. Podocytes from hypertensive and obese mice acquire an inflammatory, senescent, and aged phenotype. Am. J. Physiol. Renal Physiol. 326, F644–F660 (2024).

27. Shao, X., Xu, H., Kim, H., Ljaz, S., Beier, F., Jankowski, V., Lellig, M., Vankann, L., Werner, J. N., Chen, L., Ziegler, S., Kuppe, C., Zenke, M., Schneider, R. K., Hayat, S., Saritas, T. & Kramann, R. Generation of a conditional cellular senescence model using proximal tubule cells and fibroblasts from human kidneys. Cell Death Discov. 10, 364 (2024).

28. Liu, Y., Lv, G., Bai, J., Song, L., Ding, E., Liu, L., Tian, Y., Chen, Q., Li, K., Liu, X. & Ding, Y. Effects of HMGA2 on the epithelial-mesenchymal transition-related genes in ACHN renal cell carcinoma cells-derived xenografts in nude mice. BMC Cancer 22, 421 (2022).

29. Lin, Y.-L., Lin, Y.-W., Nhieu, J., Zhang, X. & Wei, L.-N. Sonic hedgehog-Gli1 signaling and cellular retinoic acid binding protein 1 gene regulation in motor neuron differentiation and diseases. Int. J. Mol. Sci. 21, 4125 (2020).

30. Helms, L., Marchiano, S., Stanaway, I. B., Hsiang, T.-Y., Juliar, B. A., Saini, S., Zhao, Y. T., Khanna, A., Menon, R., Alakwaa, F., Mikacenic, C., Morrell, E. D., Wurfel, M. M., Kretzler, M., Harder, J. L., Murry, C. E., Himmelfarb, J., Ruohola-Baker, H., Bhatraju, P. K., Gale, M., Jr & Freedman, B. S. Cross-validation of SARS-CoV-2 responses in kidney organoids and clinical populations. JCI Insight 6, (2021).

31. Lemos, D. R., McMurdo, M., Karaca, G., Wilflingseder, J., Leaf, I. A., Gupta, N., Miyoshi, T., Susa, K., Johnson, B. G., Soliman, K., Wang, G., Morizane, R., Bonventre, J. V. & Duffield, J. S. Interleukin-1β activates a MYC-dependent metabolic switch in kidney stromal cells necessary for progressive tubulointerstitial fibrosis. J. Am. Soc. Nephrol. 29, 1690–1705 (2018).

32. Juliar, B. A., Stanaway, I. B., Sano, F., Fu, H., Smith, K. D., Akilesh, S., Scales, S. J., El Saghir, J., Bhatraju, P. K., Liu, E., Yang, J., Lin, J., Eddy, S., Kretzler, M., Zheng, Y., Himmelfarb, J., Harder, J. L. & Freedman, B. S. Interferon-γ induces combined pyroptotic angiopathy and APOL1 expression in human kidney disease. Cell Rep. 43, 114310 (2024).

33. Mezu-Ndubuisi, O. J. & Maheshwari, A. The role of integrins in inflammation and angiogenesis. Pediatr. Res. 89, 1619–1626 (2021).

34. Cruz, N. M. & Freedman, B. S. in Methods in Cell Biology 153, 133–150 (Academic Press Inc., 2019).

35. Kim, K.-A., Wagle, M., Tran, K., Zhan, X., Dixon, M. A., Liu, S., Gros, D., Korver, W., Yonkovich, S., Tomasevic, N., Binnerts, M. & Abo, A. R-Spondin Family Members Regulate the Wnt Pathway by a Common Mechanism. Mol. Biol. Cell 19, 2588–2596 (2008).

36. Liu, J., Xiao, Q., Xiao, J., Niu, C., Li, Y., Zhang, X., Zhou, Z., Shu, G. & Yin, G. Wnt/β-catenin signalling: function, biological mechanisms, and therapeutic opportunities. Signal Transduct. Target. Ther. 7, 3 (2022).

37. Boonstra, J., Rijken, P., Humbel, B., Cremers, F., Verkleij, A. & en Henegouwen, P. van B. The epidermal growth factor. Cell Biol. Int. 19, 413–430 (1995).

38. Yu, J. S. L. & Cui, W. Proliferation, survival and metabolism: the role of PI3K/AKT/mTOR signalling in pluripotency and cell fate determination. Development 143, 3050–3060 (2016).

39. Torrens-Mas, M., Perelló-Reus, C., Navas-Enamorado, C., Ibargüen-González, L., Sanchez-Polo, A., Segura-Sampedro, J. J., Masmiquel, L., Barcelo, C. & Gonzalez-Freire, M. Organoids: An emerging tool to study aging signature across human tissues. Modeling aging with patient-derived organoids. Int. J. Mol. Sci. 22, 10547 (2021).

40. Lake, B. B., Menon, R., Winfree, S., Hu, Q., Melo Ferreira, R., Kalhor, K., Barwinska, D., Otto, E. A., Ferkowicz, M., Diep, D., Plongthongkum, N., Knoten, A., Urata, S., Mariani, L. H., Naik, A. S., Eddy, S., Zhang, B., Wu, Y., Salamon, D., Williams, J. C., Wang, X., Balderrama, K. S., Hoover, P. J., Murray, E., Marshall, J. L., Noel, T., Vijayan, A., Hartman, A., Chen, F., Waikar, S. S., Rosas, S. E., Wilson, F. P., Palevsky, P. M., Kiryluk, K., Sedor, J. R., Toto, R. D., Parikh, C. R., Kim, E. H., Satija, R., Greka, A., Macosko, E. Z., Kharchenko, P. V., Gaut, J. P., Hodgin, J. B., KPMP Consortium, Eadon, M. T., Dagher, P. C., El-Achkar, T. M., Zhang, K., Kretzler, M. & Jain, S. An atlas of healthy and injured cell states and niches in the human kidney. Nature 619, 585–594 (2023).

41. Hansen, J., Sealfon, R., Menon, R., Eadon, M. T., Lake, B. B., Steck, B., Anjani, K., Parikh, S., Sigdel, T. K., Zhang, G., Velickovic, D., Barwinska, D., Alexandrov, T., Dobi, D., Rashmi, P., Otto, E. A., Rivera, M., Rose, M. P., Anderton, C. R., Shapiro, J. P., Pamreddy, A., Winfree, S., Xiong, Y., He, Y., de Boer, I. H., Hodgin, J. B., Barisoni, L., Naik, A. S., Sharma, K., Sarwal, M. M., Zhang, K., Himmelfarb, J., Rovin, B., El-Achkar, T. M., Laszik, Z., He, J. C., Dagher, P. C., Valerius, M. T., Jain, S., Satlin, L. M., Troyanskaya, O. G., Kretzler, M., Iyengar, R., Azeloglu, E. U. & Kidney Precision Medicine Project. A reference tissue atlas for the human kidney. Sci. Adv. 8, eabn4965 (2022).

42. Petrosyan, A., Cravedi, P., Villani, V., Renieri, A., De Filippo, R., Perin, L. & Da Sacco*, S. Pd09-01 a glomerulus on a chip that recapitulates the pathophysiology of the human glomerular filtration barrier. J. Urol. 201, (2019).

43. Kroll, K. T., Homan, K. A., Uzel, S. G. M., Mata, M. M., Wolf, K. J., Rubins, J. E. & Lewis, J. A. A perfusable, vascularized kidney organoid-on-chip model. Biofabrication 16, 045003 (2024).

44. Maggiore, J. C., Ebrahimkhani, M. R. & Hukriede, N. A. A novel vascularized human kidney organoid to study podocyte and endothelial health and disease. J. Am. Soc. Nephrol. 34, 62–63 (2023).

45. Peng, K., Xie, W., Wang, T., Li, Y., de Dieu Habimana, J., Amissah, O. B., Huang, J., Chen, Y., Ni, B. & Li, Z. HIF-1α promotes kidney organoid vascularization and applications in disease modeling. Stem Cell Res. Ther. 14, 336 (2023).

46. Freedman, B. S. Physiology assays in human kidney organoids. Am. J. Physiol. Renal Physiol. 322, F625–F638 (2022).

47. Sun, X.-Y., Ju, X.-C., Li, Y., Zeng, P.-M., Wu, J., Zhou, Y.-Y., Shen, L.-B., Dong, J., Chen, Y.-J. & Luo, Z.-G. Generation of vascularized brain organoids to study neurovascular interactions. Elife 11, (2022).

48. Yousef Yengej, F. A., Pou Casellas, C., Ammerlaan, C. M. E., Olde Hanhof, C. J. A., Dilmen, E., Beumer, J., Begthel, H., Meeder, E. M. G., Hoenderop, J. G., Rookmaaker, M. B., Verhaar, M. C. & Clevers, H. Tubuloid differentiation to model the human distal nephron and collecting duct in health and disease. Cell Rep. 43, 113614 (2024).

