## Supplementary material for "Microenvironment optimization enables kidney organoid longevity with epithelial-endothelial joint basement membrane formation": Blackburnetal_Supplement

| Media Formulation | Component | Concentration |
|---|---|---|
| **Standard differentiation medium (RB)** | Advanced RPMI 1640 | 96.15% |
| | B27 | 1.92% |
| | GlutaMAX | 0.96% |
| | Penicillin/Streptomycin | 0.96% |
| **Tubuloid expansion medium (*Schutgens et al. 2019*)** | Advanced DMEM/F12 | 94.4% |
| | B27 | 1.6% |
| | R-spondin 3 conditioned medium | 1% |
| | A83-01 | 5uM |
| | FGF-10 | 100 ng/mL |
| | EGF | 50 ng/mL |
| | N-acetyl-L-cysteine | 1mM |
| | Y-27632 ROCK inhibitor | 10uM |
| | GlutaMAX | 1% |
| | HEPES buffer | 1% |
| | Penicillin/Streptomycin | 1% |
| | Primocin | 0.1 mg/mL |
| **OTEM (organoid tubular enhancing medium)** | RB | 80% |
| | Tubuloid expansion medium | 20% |
| **OVA (OTEM + VEGF + Ascorbic acid)** | OTEM | 99.8% |
| | VEGF | 1.05 uL/mL |
| | Ascorbic acid | 1.05 uL/mL |

**Supplementary Table 1. Composition of RB, OTEM, and OVA.**

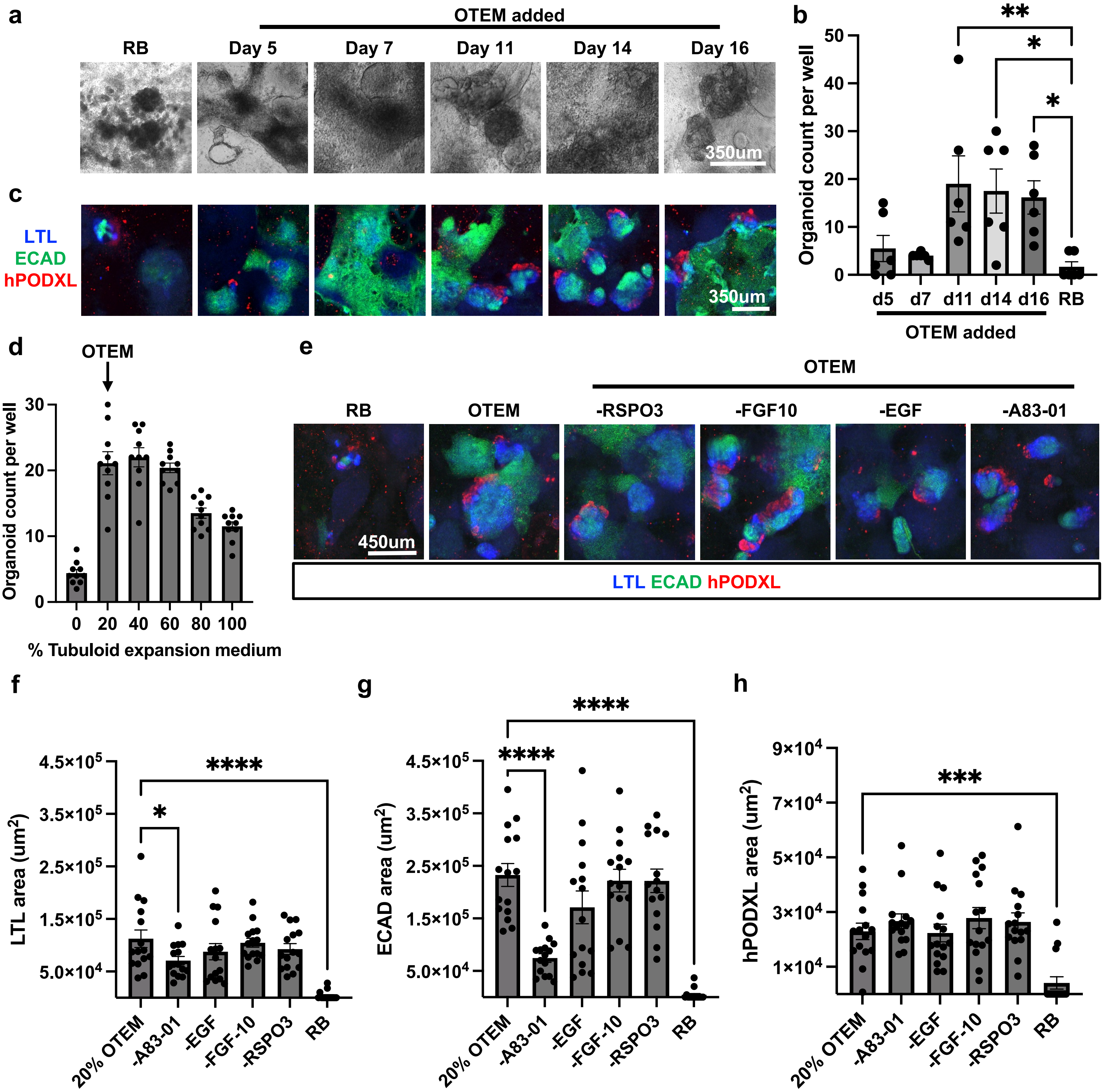


**Supplementary Figure 1. OTEM improves kidney organoid differentiation and survival. (a)** Brightfield images show changes in organoid appearance on day 31 of differentiation depending on when OTEM media is administered. Scale bars, 350um. **(b)** Number of organoids formed by day 31 depending on when OTEM is administered. Mean ± s.e.m. from n = 6 wells pooled from 3 independent experiments. One-way ANOVA performed between RB and each day of exposure. Only comparisons with $p<0.05$ are shown. *, $p<0.05$. **, $p<0.01$. **(c)** Immunofluorescence images show changes in organoid appearance on day 31 of differentiation depending on when OTEM media is administered. Scale bars, 350um. **(d)** Quantification of number of WTC11 organoids per well on day 26 of differentiation using different ratios of OTEM to RB. Mean ± s.e.m. from n = 10 wells. **(e)** IF images of representative organoids on day 31 after receiving RB, 20% OTEM, or 20% OTEM minus a single component starting on day 11 of differentiation. Scale bar, 450um. (**f-h**) Effect of media composition on (**f**) LTL, (**g**) ECAD, and (**h**) hPODXL areas on day 31. The minus conditions were fed 20% OTEM missing only the indicated growth factor. Mean ± s.e.m. from n = 15 wells pooled from 4 individual experiments. One-way ANOVA was performed between 20% OTEM and each other media composition. Only comparisons with $p<0.05$ are shown. *, $p<0.05$. ***, $p<0.001$. ****, $p<0.0001$.

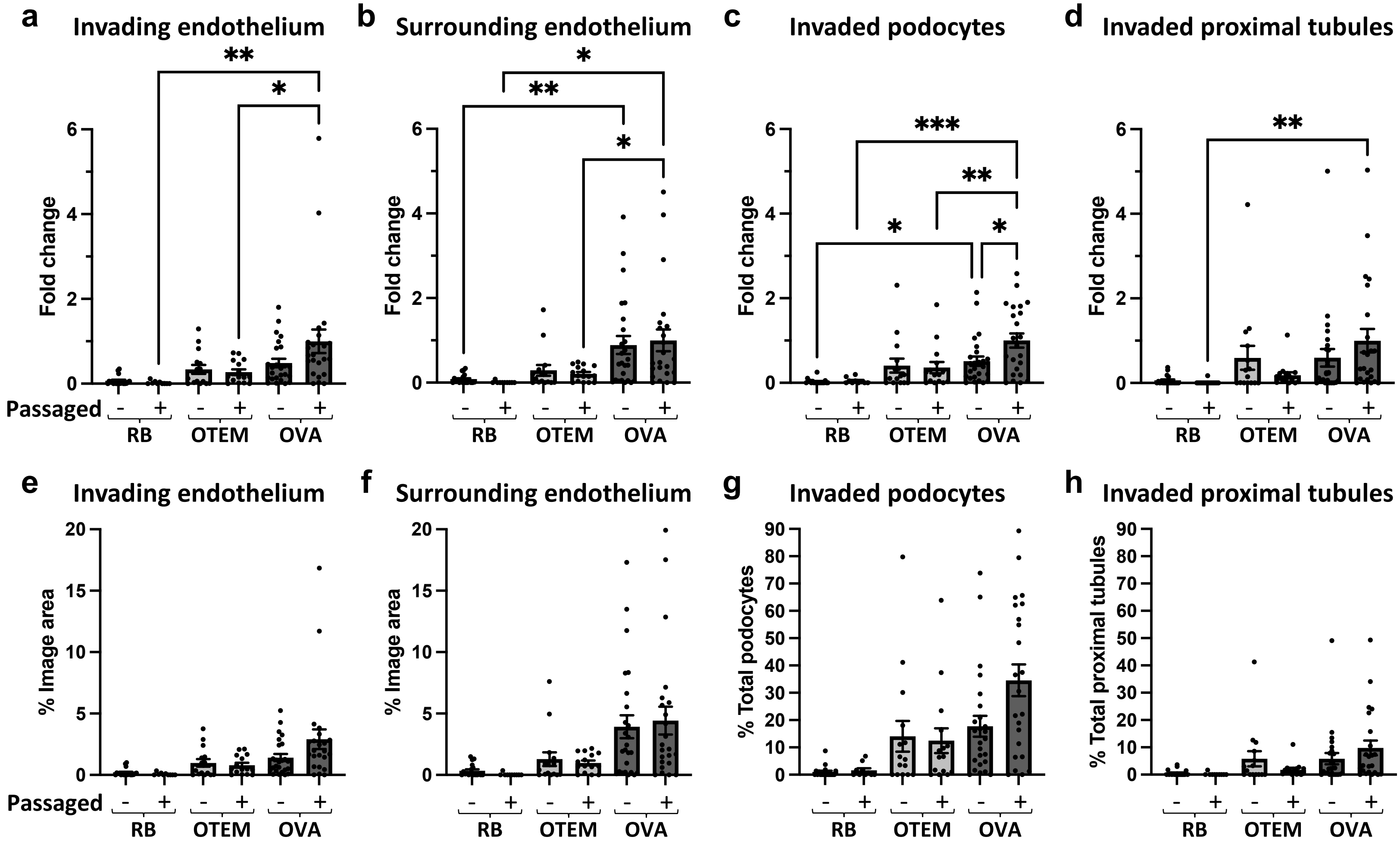


**Supplementary Figure 2. Addition of vascular factors to OTEM and passaging renal vesicles promotes endothelial differentiation and interaction with kidney organoids. (a-d)** Graphs from Figure 2 d-g with outliers included. **(e-f)** Graphs from Figure 2 d-e shown without data normalization to enable comparisons of image area. **(g-h)** Graphs from Figure 2 f-g shown without data normalization to enable comparisons of proportions of podocytes and proximal tubule invasion across conditions.

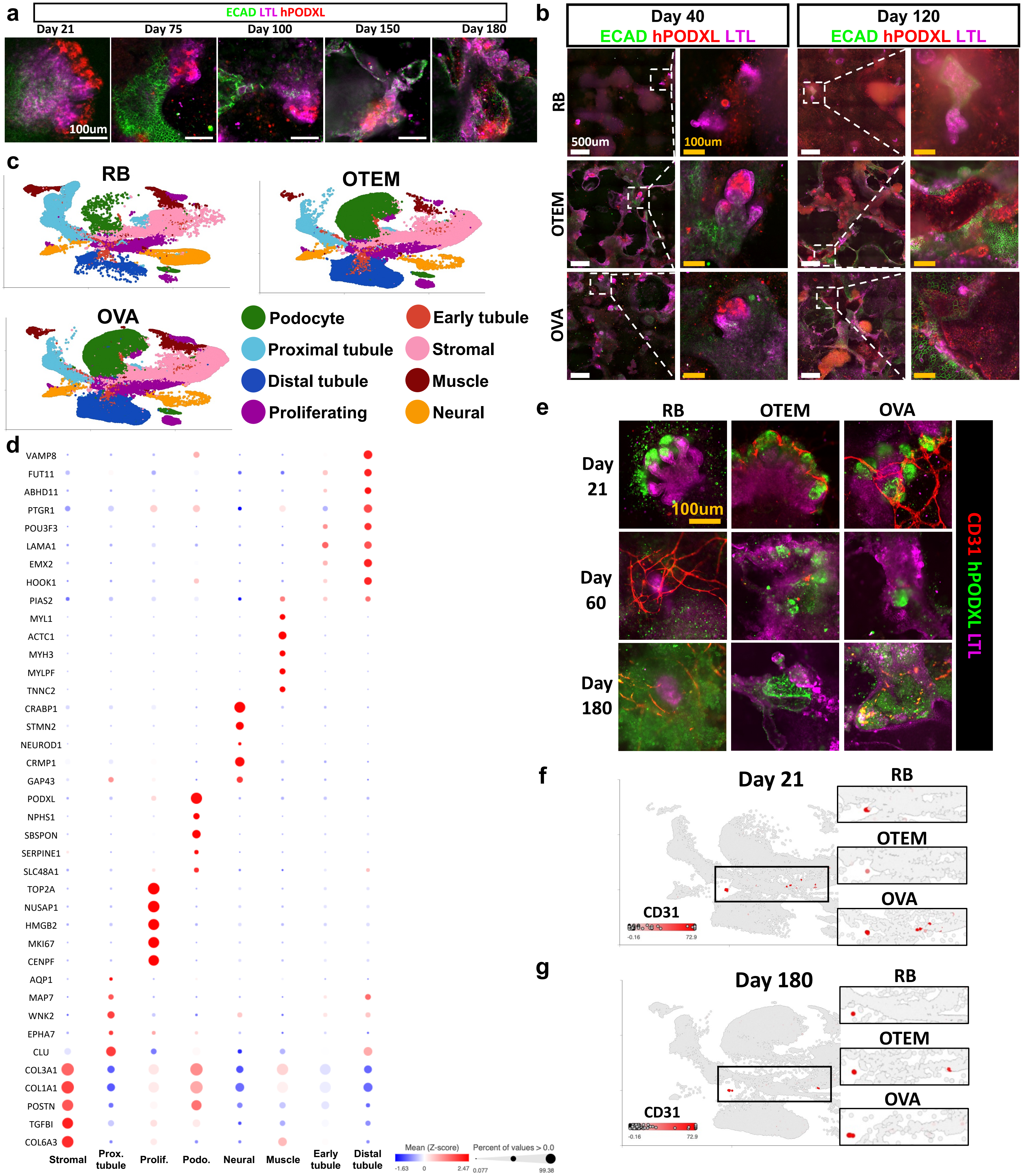


**Supplementary Figure 3. OTEM sustains kidney organoids for 6 months.** (**a**) Immunofluorescent staining from pilot experiment of kidney organoids grown in OTEM starting on day 11 and fixed at multiple timepoints up to 180 days. Scale bar 100um. (**b**) Immunofluorescent staining for classic kidney markers at days 40 and 120 in all three media conditions. These complete the time course shown in **Figure 3B.** Big picture image of organoids and surroundings has white box to indicate single organoid shown in zoomed-in image. Scale bars are 500um and 100um, respectively. (**c**) Combined 2D UMAPs showing all timepoints (Days 21, 40, 60, 120 and 180) together for each media condition. (**d**) Bubble plot showing the expression level of genes that validate the identity of each cluster from the full scRNA sequencing data set. (**e**) Representative immunofluorescent images of a single organoid stained for endothelial cells (CD31), podocytes (hPODXL), and proximal tubules (LTL) in each medium condition at days 21, 60, and 180. (**f**) Merged 2D UMAP of all media conditions at day 21 showing cells expressing CD31 in red. The region outlined by the black box is divided by media condition in the inset panels. (**g**) Merged 2D UMAP of all media conditions at day 180 showing cells expressing CD31 in red. The region outlined by the black box is divided by media condition in the inset panels.

a

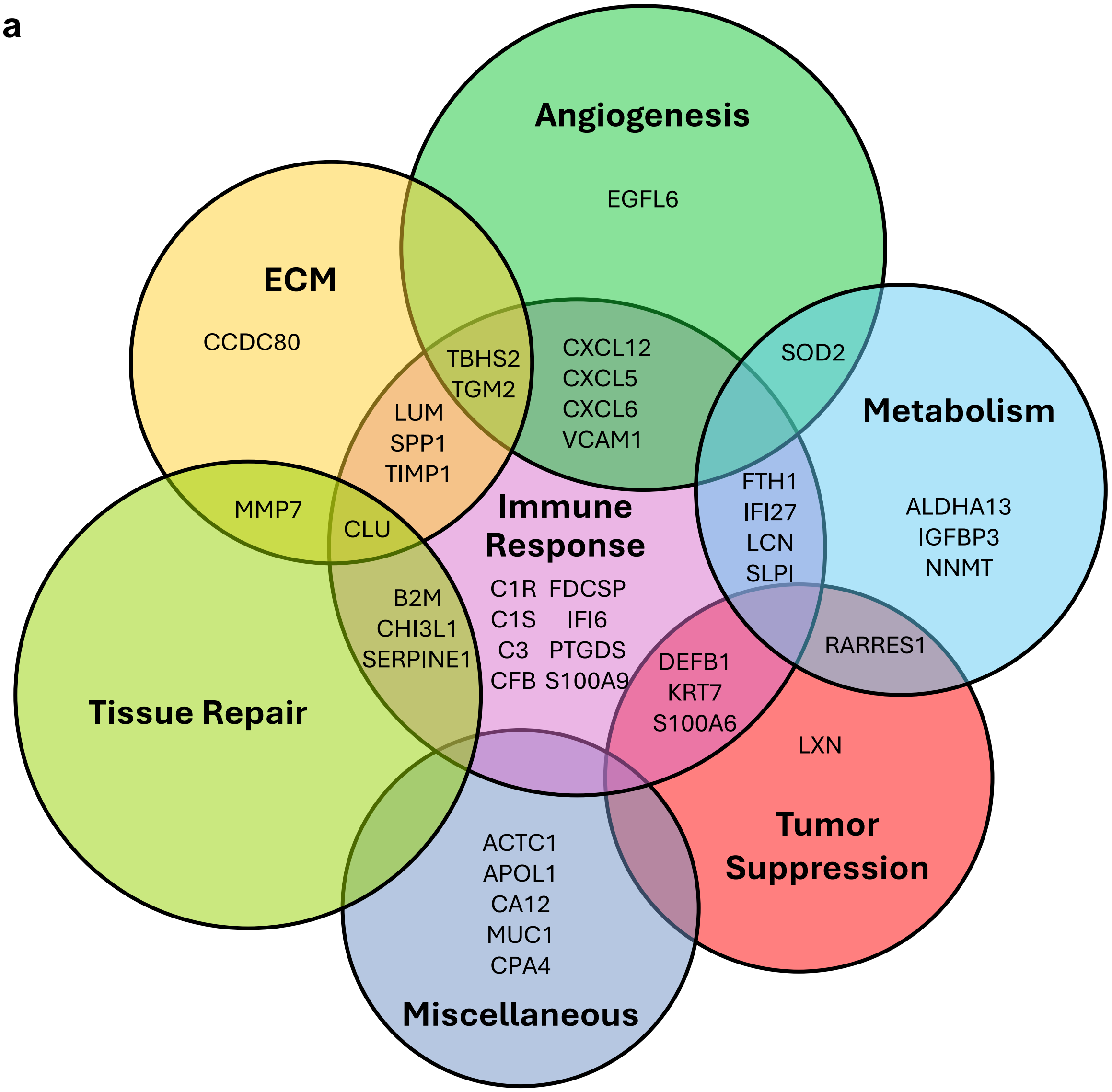


b

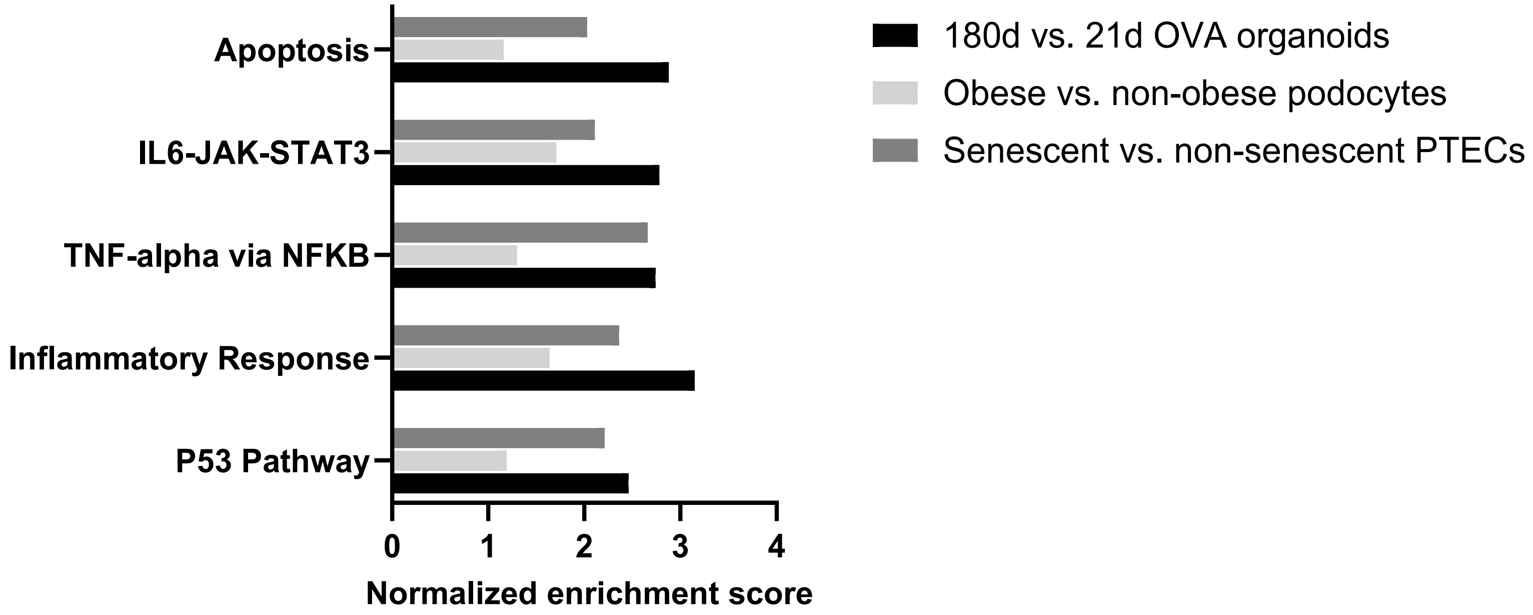


**Supplementary Figure 4. Significantly differentially expressed genes in OVA. (a)** Venn diagram of genes with Log2 fold change of at least 1 from day 21 to day 180 in OVA media. All differentially expressed genes had increased expression at day 180 v. day 21 in this condition. **(b)** Bar chart comparing normalized enrichment scores of upregulated Hallmark GSEA pathways between three independent sequencing datasets. The black bars represent 180d relative to 21d OVA-grown organoids. Light grey bars indicate murine podocytes in senescence due to obesity vs. controls. Dark grey bars indicate proximal tubule epithelial cells (PTECs) undergoing senescence after doxycycline withdrawal vs. doxycycline treated controls

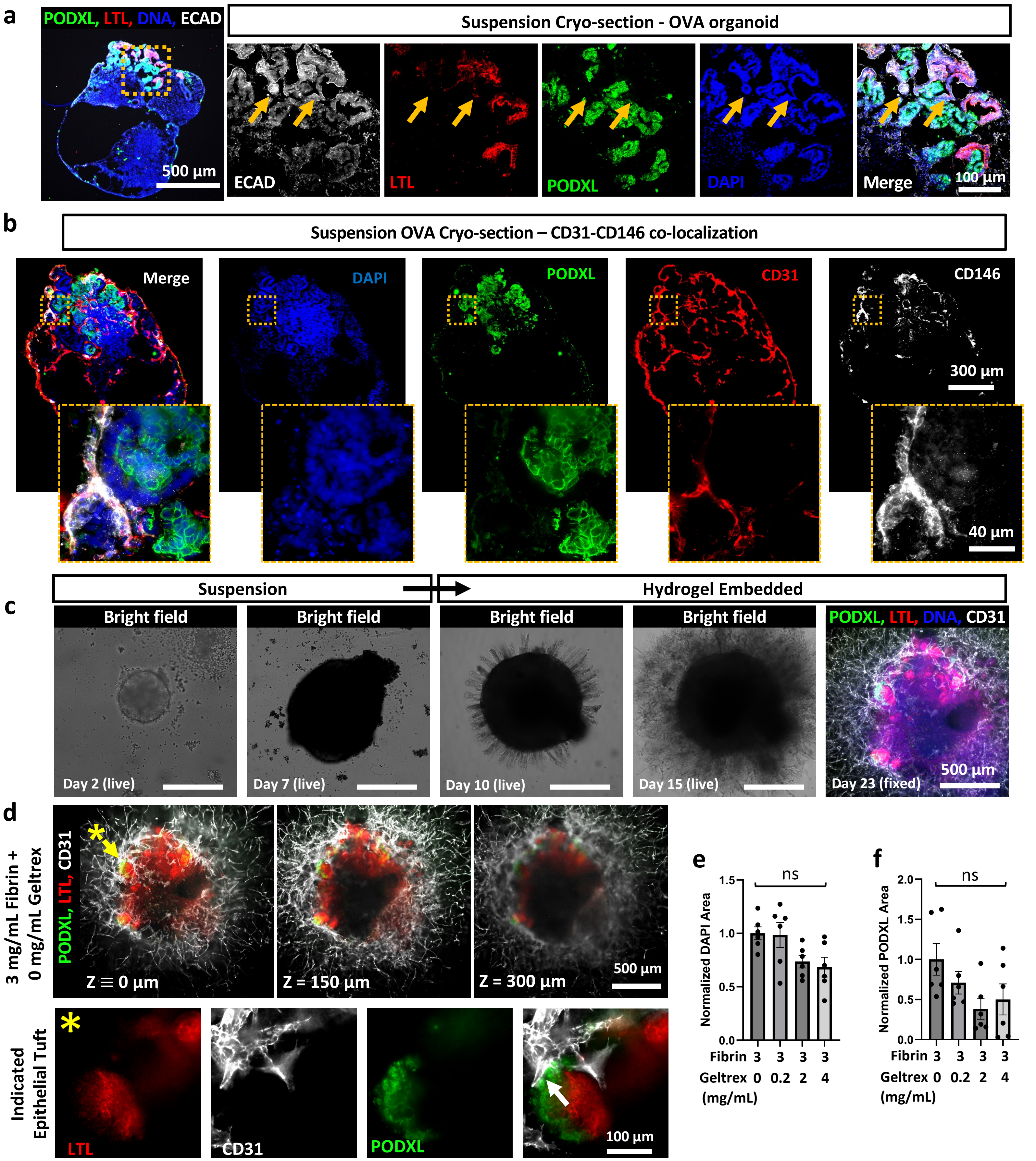


**Supplementary Figure 5. Additional characterization of suspension organoids embedded in hydrogels. (a)** Spinning disk confocal immunofluorescent images of E-cadherin (ECAD) expression in representative OVA suspension organoid cryo-section, scale bar: 500 µm. Yellow outline indicates ROI for 20x images, scale bar: 100 µm. Orange arrows indicate distal tubular ECAD+ LTL- structures. **(b)** Spinning disk confocal immunofluorescent images of CD31 and CD146 co-localization in representative OVA suspension organoid cryo-section, scale bar: 300 µm. Yellow outline indicates ROI for 60x images, scale bar: 40 µm. **(c)** Bright field time series following a suspension organoid embedded in a fibrin hydrogel and a maximum intensity projection of disk-spinning confocal z-stack after fixation. All scale bars: 500 µm. **(d)** Z-series of spinning disk confocal z-slices, at the indicated z-heights, of an organoid embedded in fibrin showing 3-dimensional emanation of the vascular network (CD31) into the surrounding hydrogel, scale bar: 500 µm. Orange asterisk (*) indicates the epithelial tuft shown at high magnification with tight association between podocytes and endothelia (white arrow), scale bar: 100 µm. **(e-f)** Immuno-fluorescence microscopy quantification of binary areas normalized to 0 mg/ml Geltrex for a given replicate showing **(e)** DAPI and **(f)** podocyte (PODXL) normalized areas per region of interest ($4mm^2$) centered on an organoid.

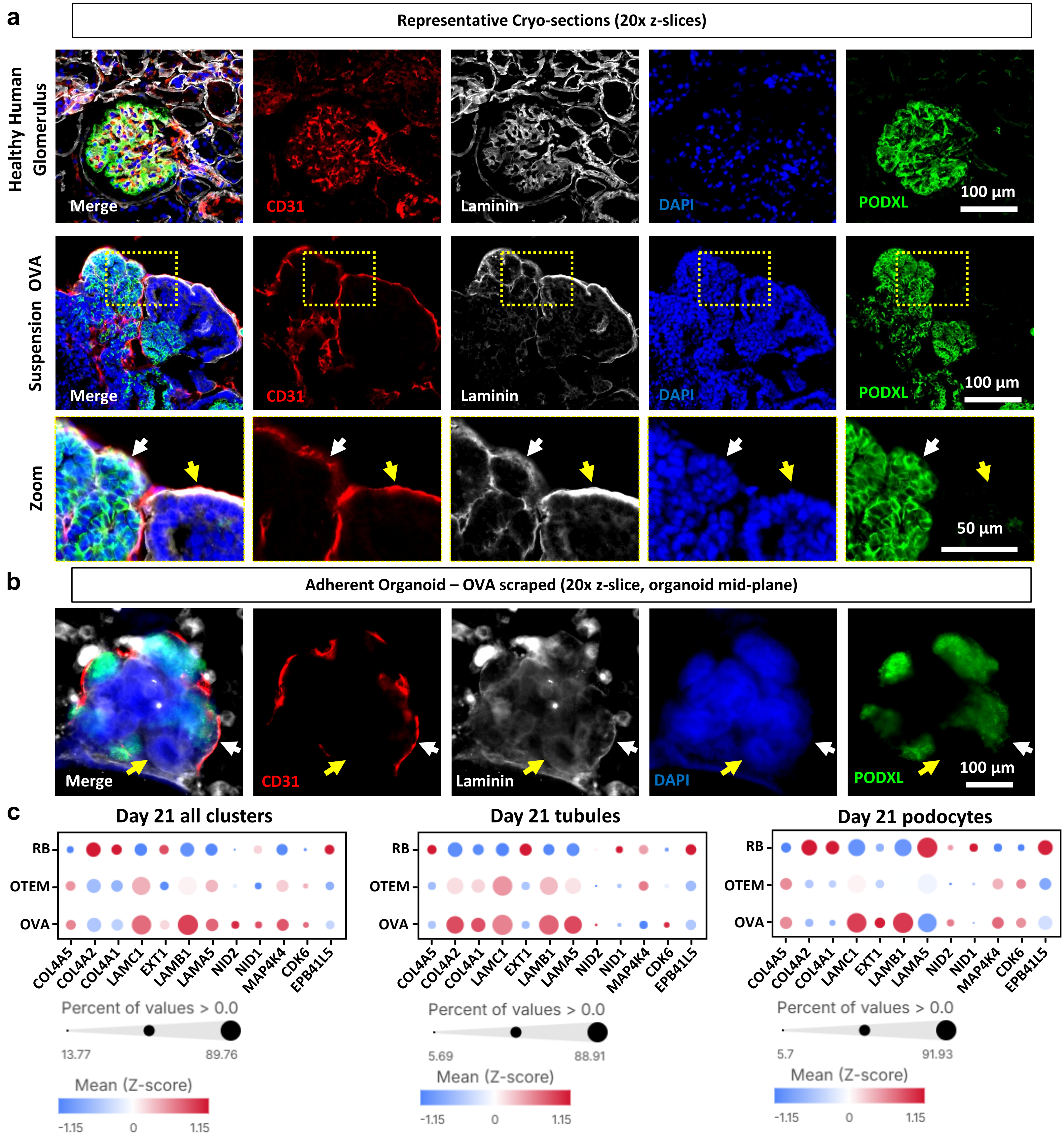


**Supplementary Figure 6. Additional characterization of organoid BM deposition and expression. (a)** Spinning disk confocal immunofluorescent images of pan-laminin staining in cryo-sections of a healthy human kidney glomerulus and representative OVA suspension organoid. 20x z-slices, scale bars: 100 µm. Yellow outline indicates ROI for digital zoom; white arrow indicates association of endothelial cells with podocyte region BM, yellow arrow indicates association with tubular BM, inset scale bar: 50 µm. **(b)** Spinning disk confocal immunofluorescent image of pan-laminin staining of a scraped and re-plated adherent organoid cultured in OVA. White arrow indicates joint BM between ECs and podocytes, yellow arrow indicates laminin expression in tubular region, scale bar: 100 µm. **(c)** Dot plot of basement membrane gene expression from scRNA-seq from all pooled clusters, tubules, and podocytes on day 21 from longevity experiments comparing adherent culture in RB, OTEM and OVA. Darker red dots indicate stronger expression across cells, and dot size reflects the percentage of cells expressing the indicated gene.
